# Measuring splash-dispersal of a major wheat pathogen in the field

**DOI:** 10.1101/2021.03.23.436423

**Authors:** Petteri Karisto, Frédéric Suffert, Alexey Mikaberidze

**Affiliations:** Plant Pathology Group, Institute of Integrative Biology, ETH Zurich, Zurich, Switzerland; Université Paris-Saclay, INRAE, AgroParisTech, UMR BIOGER, 78850 Thiverval-Grignon, France

**Keywords:** Dispersal kernel, Dispersal gradient, *Zymoseptoria tritici*, Wheat, Rain-splash dispersal, Bootstrap, Spatially-explicit modeling, Plant disease epidemiology, Dispersal ecology

## Abstract

Capacity for dispersal is a fundamental fitness component of plant pathogens. Empirical characterization of plant pathogen dispersal is of prime importance for understanding how plant pathogen populations change in time and space. We measured dispersal of *Zymoseptoria tritici* in natural environment. Primary disease gradients were produced by rain-splash driven dispersal and subsequent transmission via asexual pycnidiospores from infected source. To achieve this, we inoculated field plots of wheat (*Triticum aestivum*) with two distinct *Z. tritici* strains and a 50/50 mixture of the two strains. We measured effective dispersal of the *Z. tritici* population based on pycnidia counts using automated image analysis. The data were analyzed using a spatially-explicit mathematical model that takes into account the spatial extent of the source. We employed robust bootstrapping methods for statistical testing and adopted a two-dimensional hypotheses test based on the kernel density estimation of the bootstrap distribution of parameter values. Genotyping of re-isolated pathogen strains with strain-specific PCR-reaction further confirmed the conclusions drawn from the phenotypic data. The methodology presented here can be applied to other plant pathosystems.

We achieved the first estimates of the dispersal kernel of the pathogen in field conditions. The characteristic spatial scale of dispersal is tens of centimeters – consistent with previous studies in controlled conditions. Our estimation of the dispersal kernel can be used to parameterize epidemiological models that describe spatial-temporal disease dynamics within individual wheat fields. The results have the potential to inform spatially targeted control of crop diseases in the context of precision agriculture.

## Introduction

Ability to spread within host populations is a fundamental requirement for plant pathogens. Spatial spread directly influences the number of new hosts plants that a pathogen can potentially invade and affects the spatial distribution of the pathogen population. For polycyclic diseases, even small differences in the spread during one infection cycle can result in considerable differences in epidemic outcomes after several cycles. Understanding the mechanisms and scale of the spread improves our ability to predict and control plant disease epidemics.

Although the epidemiological importance of dispersal has long been recognized (Heald, 1913), measurements in field conditions are rare, for numerous reasons: (i) field experiments on dispersal are difficult to design and to conduct (McCartney et al., 2006). (ii) They are inherently multidisciplinary spanning the interface between biology and physics, (iii) It is challenging to measure the primary dispersal gradient (i.e., resulting from a single round of pathogen reproduction) independently of secondary dispersal and environmental gradients (Gregory, 1968). (iv) Characterization of finer scale dispersal occurring on sub-meter scales can be difficult in the field. Existing literature contains many more studies characterizing dispersal of airborne dispersal of plant pathogens compared to splash dispersal (McCartney et al., 2006): Fitt et al. (1987) found 305 datasets on air-borne dispersal in the field, only 10 datasets on splash-dispersal in controlled conditions, and no studies on splash-dispersal in the field. A possible explanation for this disparity is that commercially available spore traps or disease assessments readily capture spatial scales of meters or longer, characteristic of air-borne dispersal, but those are more challenging to apply over sub-meter scales typical for splash-dispersal.

*Zymoseptoria tritici* (formerly *Mycosphaerella graminicola)* is a major fungal pathogen of wheat (Jørgensen et al., 2014; Dean et al., 2012) that causes septoria tritici blotch (STB) disease. *Z. tritici* disperses by means of air-borne ascospores (sexual) and splash-borne pycnidiospores (asexual). Pycnidiospores are the main driver of polycyclic epidemics during the wheat growing season. Ascospores are the main source of primary inoculum in the beginning of an epidemic and contribute to epidemic development during the season (Zhan et al., 1998, 2000; Duvivier et al., 2013). According to some studies, initial inoculum via air-borne ascospores is uniform across the horizontal spatial extent of wheat fields and does not represent a limiting factor for epidemic development (Morais et al., 2016), implying that vertical dispersal from lower to higher leaf layers is more relevant epidemiologically than horizontal dispersal across the spatial extent of wheat fields. However, it is not clear how general these conclusions are: it is possible that they hold only in regions with temperate wet climate and intense wheat production dominated by STB-susceptible wheat cultivars, where the studies have been conducted. Yet most of research on splash dispersal of *Z. tritici* has focused on vertical dispersal of spores from initial infection on basal leaves of seedlings to emerging leaf layers (Shaw, 1987; Lovell et al., 1997; Bannon and Cooke, 1998; Lovell et al., 2004*b*; Vidal et al., 2018). In this context, the interaction between the pathogen and its host plant has been described as a “race”, where the pathogen population needs to “climb” up to the next leaf layer before current layer becomes senescent and resources are depleted (Robert et al., 2018).

We argue that horizontal dispersal is as important as vertical dispersal, because it greatly influences the ability of a specific pathogen genotype to take over a field and therefore it can play a major role in the dynamics of emerging pathogen genotypes adapted to control measures such as fungicides or disease resistance genes in host plant. However, because of insufficient empirical knowledge on rain-splash driven horizontal dispersal, it has not been considered in modeling the emergence of new pathogen strains (e.g.,(Mikaberidze et al., 2017; Willocquet et al., 2020)). Horizontal expansion of disease foci (Zadoks and van den Bosch, 1994) corresponds to the growth of a genetically uniform pathogen population which is well-adapted control measures applied in a spatially uniform manner (e.g., fungicides or disease resistance genes). For this reason, spatially heterogeneous control strategies have been suggested such as varietal mixtures in individual fields or landscape mosaics (Mundt and Browning, 1985; Brophy and Mundt, 1991; Newton et al., 2009; Sapoukhina et al., 2010; Newton and Guy, 2011; Mikaberidze et al., 2014; Djidjou-Demasse et al., 2017; Ben M’Barek et al., 2020). Spatial scale of such strategies can be optimized based on the knowledge on spatial scales of horizontal dispersal of the pathogen. This knowledge is captured by a dispersal kernel function that describes the probability of an individual dispersal event to end up at a certain location relative to the source (Nathan et al., 2012).

Dispersal via asexual spores of *Z. tritici* and of the wheat pathogen *Parastagonospora nodorum* (formerly *Septoria nodorum)* has been studied in controlled conditions using inoculations via infected straw or spore suspension combined with artificial rain (Brennan et al., 1985; Saint-Jean et al., 2004; Vidal et al., 2017). Bannon and Cooke (1998) studied the effect of wheat-clover intercrop on dispersal from plates by artificial rain and detected a reduction of dispersed spores at 15 cm distance from the source. No experiment has been conducted within a host plant canopy in the field that would allow to parameterize the dispersal kernels associated with splash dispersal.

We conducted an experiment to measure splash-dispersal of *Z. tritici* in field conditions. We carried out localized artificial inoculations with two *Z. tritici* strains as well as their mixture that contained equal proportions of each strain. Conducive weather conditions allowed us to measure primary dispersal gradients resulting from single dispersal events following the initial artificial inoculation. We used automated digital image analysis (Karisto et al., 2018) to estimate sizes of pathogen populations on wheat leaves based on detection of pycnidia (fungal fruiting bodies). In addition, we genotyped the re-isolated strains using strain-specific primers to differentiate the inoculated pathogen populations from the background natural population. Using a spatially-explicit mathematical model that took into account the spatial extent of the source area (Karisto et al., 2019*b*), we achieved the first estimates of dispersal kernel parameters associated with splash dispersal of *Z. tritici* pycnidiospores in field conditions.

## Materials and methods

### Plant materials and agronomic practices

The experiment was performed at the Field Phenotyping Platform site, Eschikon Field Station, ETH Zurich, Switzerland (Kirchgessner et al., 2017). Experimental plots were sown with winter wheat (*Triticum aestivum)* cultivar Runal (Breeder: Swiss Federal Research Station for Agroecology and Agriculture, FAL, Zurich, Switzerland) on 1 November 2016. Sowing density was 440 seeds/m^2^ and the observed stem density on 19 June 2017 was 730 stems/m^2^. Field maintenance included herbicide Herold SC (0.6 l/ha; Bayer) on 2 November 2016, and stem shortener Moddus (0.5 l/ha; Syngenta) on 13 April 2017. Fungicide Input (1.25 l/ha; Spiroxamin 300 g/l, Prothioconazol 160 g/l; Bayer) was applied on 13 March 2017 to suppress the natural epidemic of STB and other fungal diseases.

Similar experiment was also prepared at INRAE Bioger, Thiverval-Grignon, France (coordinates: 48.84ON, l.952E). The experimental design was slightly adapted. Due to unconducive weather conditions the inoculation did not produce measurable primary disease gradients. Therefore, no data is presented.

### Experimental design

The experimental plots were 1.125m × 4m rectangles consisting of nine 4m-long rows with the distance of 12.5 cm between the rows. Plots were randomly assigned to four treatments with five replicates of each treatment as shown in Fig. 1A. The four treatments were: (i) inoculation with the single strain ST99CH_1A5 (short identifier 1A5), (ii) inoculation with the single strain ST99CH_3D7 (short identifier 3D7), (iii) inoculation with both strains 1A5 and 3D7 (mixed inoculation) and (iv) control (no inoculation). The two strains were collected and isolated in Switzerland in 1999 (Zhan et al., 2002, see also www.septoria-tritici-blotch.net/isolate-collections.html). The strains were chosen because of their capacity to infect cultivar Runal, their contrasting production of pycnidia (Stewart et al., 2018), and their ability to cross with each other (1A5, Ma1-1; 3D7, Mat 1-2: (Suffert et al., 2018)).

**Figure 1:**
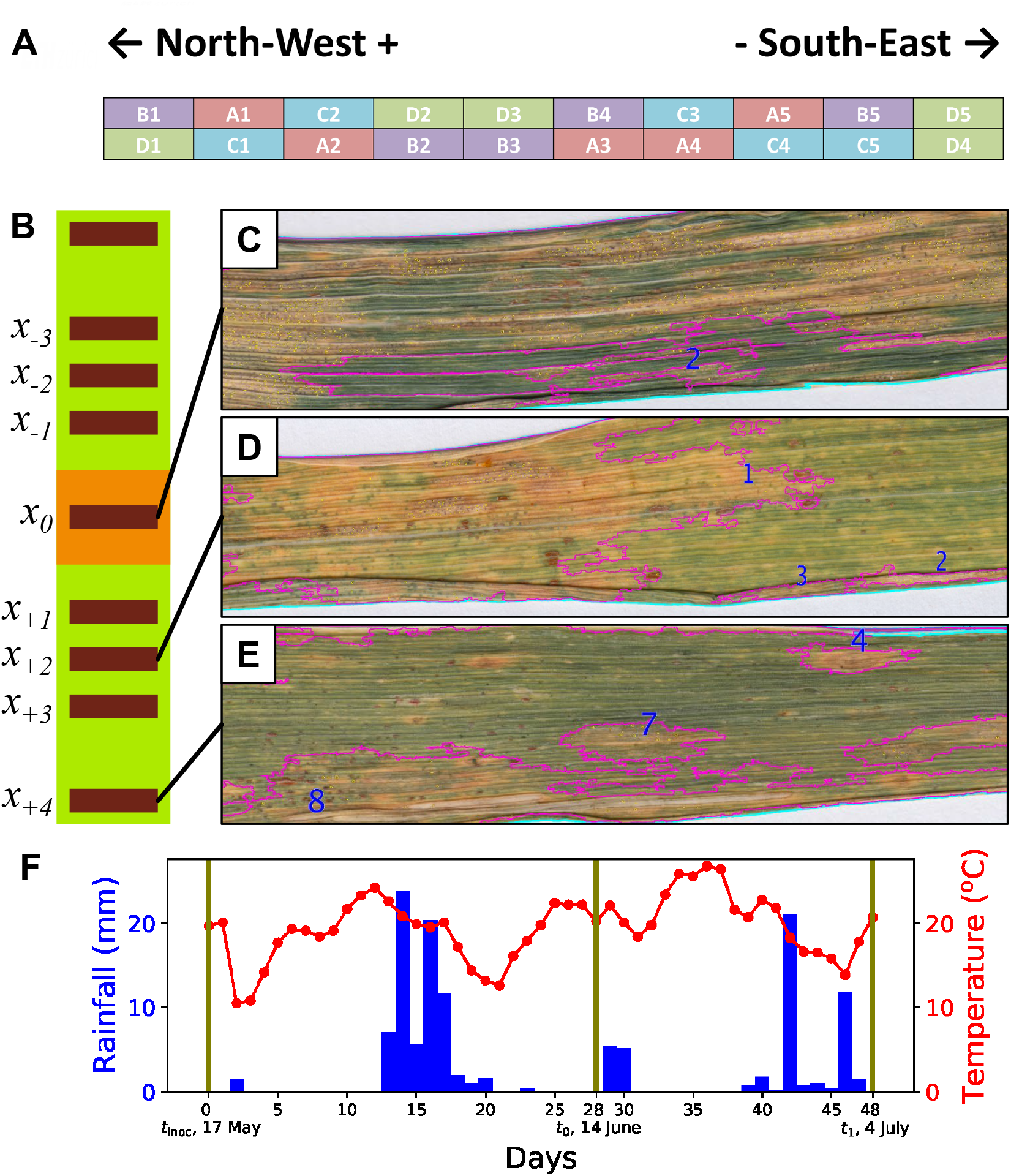
Overview of the experiment. (A) Arrangement of the plots in the field (treatment letter and replicate number: A = 1A5, B = mixed inoculation, C = control, D = 3D7.) (B) Design of an experimental plot. 40 cm-wide inoculation area in the middle (orange). Distances from the middle of the inoculated area to the middle of each measurement line were *x*_0_ = 0 cm, *x*_±1_ = 40 cm, *x*_±2_ = 60 cm, *x*_±3_ = 80 cm and *x*_±4_ = 120 cm. (C, D, E) Overlay images illustrate the automated image analysis by showing leaves collected from measurement lines *x*_0_ (C), *x*_+2_ (D) and *x*_+4_ (E) of the treatment 3D7 at the sampling date *t*_1_. Cyan, purple and yellow lines mark the borders of leaves, lesions and pycnidia, respectively. (F) Weather conditions during the experiment. Daily precipitation in the blue bars, daily mean temperature in the red line, the dates of the inoculation and the leaf samplings are shown with vertical lines.

In each plot, a 40 cm-wide area across the middle of the plot was inoculated. Disease levels were measured in the middle of the inoculated area (*x*_0_ = 0 cm) and at eight locations outside of the inoculated area, four on each side, at distances *x*_±1_ = 40 cm, *x*_±2_ = 60 cm, *x*_±3_ = 80 cm and *x*_±4_ = 120 cm from the center of the inoculated area (see Fig. 1B). The areas in which we conducted measurements (“measurement lines”, shown as dark rectangles in Fig. 1B) were 10 cm wide and 87.5 cm long across the plot excluding 12.5 cm borders at each edge of the plot to reduce edge effects. Disease assessments were conducted uniformly over the area of each measurement line.

### *Z. tritici* inoculation

Inoculum was prepared by growing the fungus for seven days in yeast-sucrose-broth (https://dx.doi.org/10.17504/protocols.io.mctc2wn). The liquid culture was then filtered, the blastospores were pelleted in centrifuge and re-suspended into sterile water. The washed spore suspension was diluted with water to achieve the concentration of 10^6^ spores/ml. For mixed inoculation, the spore suspension was obtained by mixing the same volume of each single strain suspension so that the final spore concentration was 10^6^ spores/ml and each strain was present with the concentration of 5 × 10^5^ spores/ml. Finally, we added 0.1 % (v/v) of Tween20 and kept the inoculum suspension on ice until spraying on the same days.

Inoculation was performed by spraying 300 ml of spore suspension onto the inoculation site of each plot using a hand-pump pressurized sprayer. The plots were inoculated during the late afternoon to avoid direct sunlight. All treatments were inoculated with the same sprayer, which was rinsed with water and with 70% ethanol to clean all parts before inoculating each treatment. Entire canopy within the inoculation area was inoculated until runoff. During the spraying, the 40 cm-wide inoculation area was bordered with plastic sheets to avoid spillover of the inoculum outside the area. After spraying, the border sheets were folded over the canopy to enclose the plants in a plastic tent to maintain high humidity overnight. The tents were removed early next morning to avoid overheating. The inoculation was repeated in the next evening. Photographs documenting the inoculation process are shown in Appendix B.

The first attempt to inoculate was made on 5 and 6 April, when F-3 layer (the third leaf layer below flag leaf) had mostly emerged (approximate growth stage, GS 22, Zadoks et al. (1974)), and inoculation success was assessed on 24 April and again on 3 May. Due to cold weather the infection was extremely limited: we observed low levels of disease in the F-3 leaf layer when the plants were in the beginning of stem elongation (F-1 emerging, GS 35). Average STB incidence in F-3 layer in the inoculated area on 3 May was 6.1%,4.9% 2.9%, and 0% for strains 1A5, 3D7, mixed inoculation and control, respectively. We considered this inoculation as failed for the purpose of the experiment, because the secondary spread from such low initial infection levels would likely cause only negligible disease gradients. We decided to inoculate again the higher leaf layers to achieve stronger, measurable disease gradients. Dates of this second, main inoculation were 17 and 18 May 2017, when flag leaves have already emerged (GS 39-41).

### Assessment of disease gradients

The disease assessment combined incidence and severity measurements. It was performed at two time points. At *t*_0_, on 14 June 2017 (GS 70) only the inoculation areas were assessed carefully to evaluate the success of inoculation across the measurement line *x*_0_. Flag leaves outside the inoculation area were inspected visually and confirmed to be healthy. At *t*_1_ on 4 July 2017 (GS 85) all measurement (*x*_0_, *x*_±1_, *x*_±2_, *x*_±3_ and *x*_±4_) lines were assessed in all plots corresponding to inoculated treatments (1A5, 3D7 and mixed inoculation), while only one randomly chosen measurement line was assessed in each of the control plots.

At *t*_0_, STB incidence was measured at the leaf scale in the following manner. Thirty to forty stems were inspected on each measurement line. The highest diseased leaf layer was recorded for each stem. The leaves lower than that were assumed to be diseased as STB is usually more prevalent in the lower leaf layers (Lovell et al., 2004*a*). Additionally, naturally senescent leaf layers at the bottom of the canopy were recorded. In this way, incidence was estimated for all non-senescent leaf layers. After estimating the incidence, eight infected leaves were collected from up to two consecutive leaf layers that had incidence higher than 20% (to avoid removing too much of the inoculum). The collected leaves were then mounted on paper sheets and scanned with 1200 dpi resolution. The resulting images were analyzed using automated image analysis method to measure two aspects of conditional STB intensity that represent the host damage and the pathogen reproduction, as described in Karisto et al. (2018). Host damage was measured as the percentage of leaf area covered by lesions and pathogen reproduction was measured as the number of pycnidia per leaf. The sampled leaf layers at *t*_0_ were the flag leaf layer (F) and the layer immediately below the flag leaf (F-1).

At *t*_1_, the wheat leaves were already mostly chlorotic and hence the incidence measurement was not possible in the field. Instead, we collected at random about 24 leaves from each measurement line. The leaves were taken into the lab and inspected for the presence of pycnidia. Incidence was recorded based on the presence of pycnidia, and the leaves with pycnidia were scanned as described above to quantify the conditional STB intensity. Due to vast chlorosis, the measurement of host damage was considered unreliable and only pathogen reproduction was used in the subsequent analysis. Thus, we measured the conditional disease intensity using only the numbers of pycnidia per leaf.

We estimated the number of asexual reproduction and dispersal events between *t*_0_ and *t*_1_ using the following arguments. First, based on the data from Shaw (1990, as revisited in Fig. A1 of Karisto et al. (2018)), the latent period after inoculation was estimated to be longer than 20 days (average daily temperature during first 19 days was 19°C). Thus, there was likely no spread from the inoculation areas during the rainy period only 13-17 days after inoculation (dai) (Fig. 1F). This was further confirmed by a visual assessment on 8 June (22dai), when we observed only few tiny lesions and mostly no pycnidia on plants inside the inoculation areas, concluding that substantial spread did not yet occur. Second, at *t*_0_ (14 June, 28 dai) we observed substantial disease (Fig. 2A) in the inoculation areas. There was no rain during the week preceding *t_0_,* two strong showers during the night about 36 hours after *t*_0_ and no more rain during the following week. We conclude that there was most likely only one asexual spread event right after *t*_0_, which caused the disease gradients that we observed at *t*_1_ (48 dai) outside of the inoculation areas.

**Figure 2:**
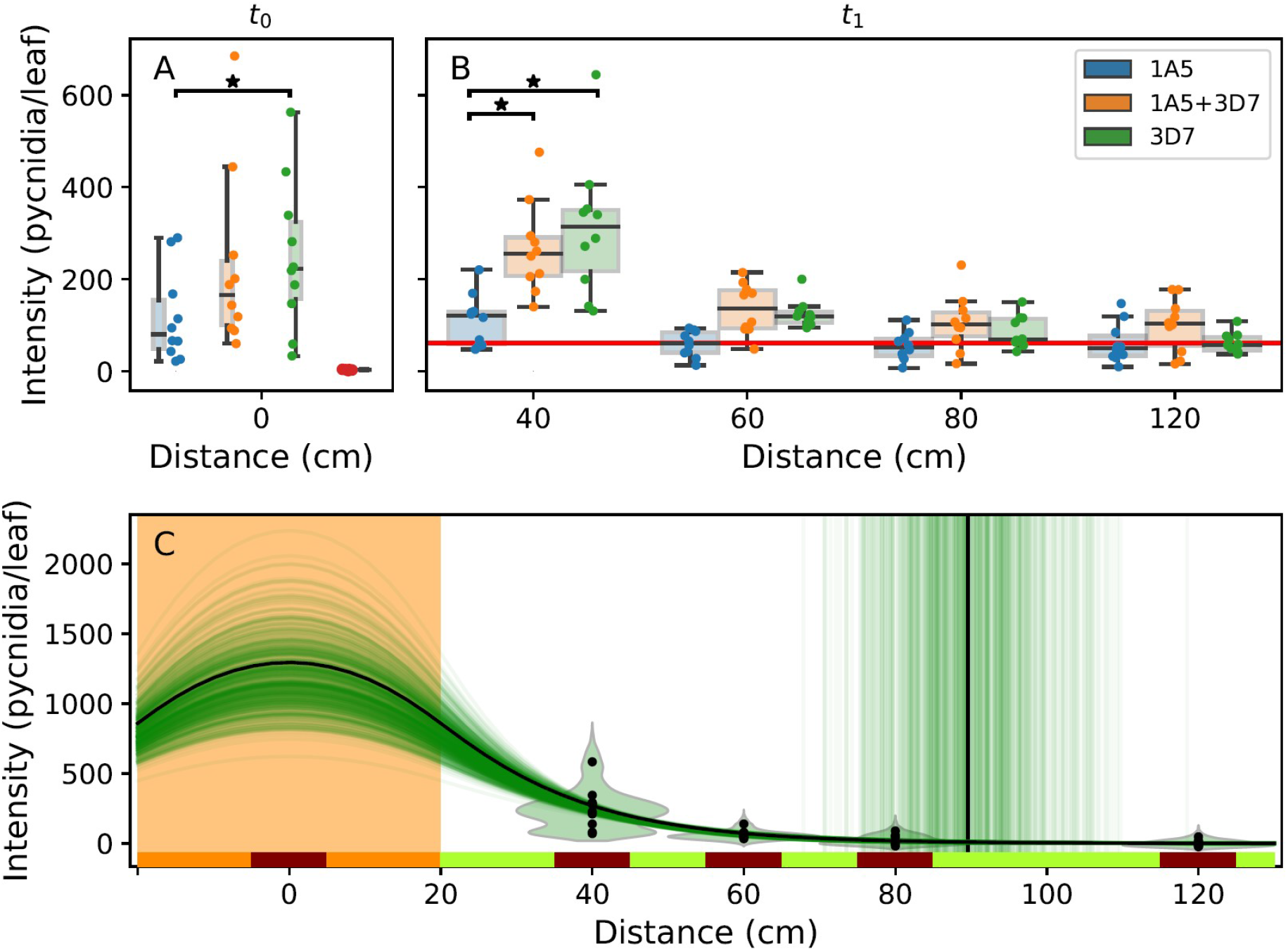
Disease intensity and dispersal gradients. In all panels, dots represent means over individual replicate plots. In panels A and B, we present outcomes for 1A5 (blue), mixed inoculation 1A5—3D7 (orange), 3D7 (green), and non-inoculated control (red) treatments. Black horizontal bars with asterisks show the significant pairwise differences between treatments at *t*_0_ and at *t*_1_. A: Disease intensity in the inoculation area at *t*_0_, measurements for leaf layers F and F-1 are shown for the three treatments, while layer F-2 for control (as there was no disease at higher leaf layers). B: Gradients of disease intensity in leaf layer F at *t*_1_ after the spread event for the three inoculation treatments. Red horizontal line shows the mean disease intensity in F layer over all control plots. C: Disease gradients of strain 3D7 are fitted using Eq. 3. Black curve shows the best fit, vertical black line shows 99^th^ percentile of the dispersal distance (*x*_99_). Green violin plots show the distributions of 500 bootstrap replicates, green curves represent fits to each of these bootstrap replicates and green vertical lines show the corresponding *x*_99_-values.

Based on the arguments above, we conclude that the second inoculation was successful: it caused high disease levels observed in the inoculation areas at *t*_0_, after a latent period of 3-4 weeks. After three more weeks, at *t*_1_, there were clearly visible symptoms outside of the inoculation areas. The observed symptoms at *t*_1_ can be entirely attributed to the rain event on 17 June (shortly right after *t*_0_) and consequent asexual spread of the pathogen.

### Genotyping

To confirm that the observed disease gradients were due to the experimental treatments, we re-isolated 153 *Z. tritici* individuals from individual pycnidia on the collected leaves after scanning. The isolations were performed from leaves collected in the measurement lines *x*_±1_ and *x*_±3_. To detect the inoculated strains, we designed and used strain-specific PCR-primers (see details in the Appendix A.). As a result of the genotyping, the isolates were identified as either 1A5 or 3D7 if the two primer pairs specific to one of these strains yielded amplification, while the primer pairs specific to the other strain did not.

### Statistical analysis

#### Fitting disease gradients

The disease intensity (numbers of pycnidia per leaf) at *t*_1_ in a given measurement line is a result of dispersal of asexual spores from the inoculation area and successful infections within the measurement line. Assuming spatially uniform success of infections in all plots, the observed disease gradient reflects the dispersal gradient of spores and thus provides the effective dispersal gradient of the pathogen population. Following the spatially-explicit method for parameterizing a dispersal kernel (Karisto et al., 2019*b*), the dispersal process can be described mathematically using two area integrals: one over the source area and the other over the destination area. We used the exponential dispersal kernel that fits well for splash dispersal (Fitt et al., 1987; Saint-Jean et al., 2004).

The disease intensity at *t*_1_ in a measurement line at distance *x*\* from the inoculation area is given by

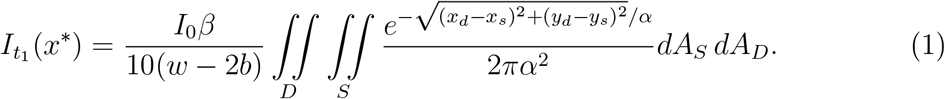

This general approach can take into account arbitrary shapes of the two areas of integration D and *S.* In the specific case of our experimental design, the two areas correspond to the following rectangles: *D* = {(*x_d_,y_d_*)} = [*x*\* — 5cm,*x*\* + 5cm] × [*b,w* − *b*] is the destination area that corresponds to a measurement line (the dark rectangles in Fig. 1B), *S* = {(*x_s_,y_s_*)} = [0, *l_s_*] × [0, *w*] is the source area that corresponds to the inoculated area (the orange rectangle in Fig 1B). *I_0_* is the STB intensity (numbers of pycnidia per leaf) in the inoculation area at *t_0_; β* is the transmission rate (unitless) describing the number of new pycnidia produced in the measured leaf layer per unit of measured intensity in the source leaves; *w* = 112.5 cm is the plot width; *l_s_ =* 40 cm is the extent of the inoculated area along the *x*-axis (i.e., the length of the plot); *b* = 12.5cm is the width of the border excluded from the measurement lines; and *α* is the shape parameter of the dispersal kernel that determines the characteristic spatial scale of dispersal (below we call it the “dispersal parameter”). The measurement lines were 10 cm-wide. Hence, the integration over a measurement line divided by the area of the line, 10 cm × (*w* ‒ 2*b*), gives the average disease intensity in the measurement line, reflecting spatially uniform leaf sampling.

Note that 10 cm width of the measurement lines was practically the smallest possible width that could be achieved in the field measurements, because the foliage of even a single straw spans more than 10 cm, limiting the spatial resolution of our measurements. For this reason, we simplified the model by neglecting the width of the measurement lines and assigning the disease intensity values of each measurement line to the middle of the line. With this simplification, the disease gradient is calculated as

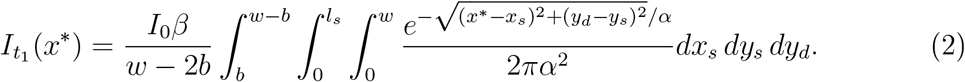

As implied by Madden et al. (2007) and the analysis of Fitt et al. (1987), dispersal is often modeled as a one-dimensional process. This simplification may lead to incorrect estimates of the dispersal kernel (Karisto et al., 2019*b*). However, to enable comparisons between our outcomes and the earlier estimates in controlled conditions, and to highlight the distorting effect of this “standard” simplification, we constructed the one-dimensional function describing the dispersal process that follows an exponential kernel:

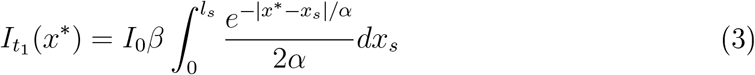

In Eq. (3) the integral takes sum over the inoculated area along the length of the plot.

We conducted the integration in Eq. (3) and obtained a simpler expression:

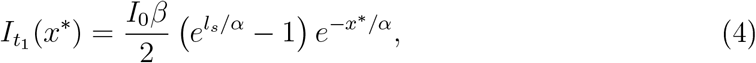

where *x*\* takes the values outside of the inoculation area.

#### Estimation of dispersal and transmission parameters

The data used in the analysis were obtained in the following way. First, we conducted STB incidence measurements and we acquired conditional STB intensity measurements from the digital image analysis. Second, the STB intensity values in each measurement line were multiplied by the corresponding STB incidence to obtain the full or “unconditional” intensity values, thereafter called “intensity” for the sake of simplicity. Third, we calculated the average full intensity in each measurement line to obtain five data points (from five replicates) for in each treatment (at distances *x*_0_, *x*_±1_, *x*_±2_, *x*_±3_ and *x*_±4_). These average values over each measurement line were used for fitting the model functions. The dispersal gradient functions (Eqs. (1), (2)) were fitted to the data to estimate *α* and *I*_0_*β*.

To compare the treatments and the two directions, we used the bootstrapping approach. We re-sampled the collected samples with replacement to create a large set of bootstrap samples. Variation in the bootstrap samples reflects the variation that we expect to observe if the actual experiment was repeated several times (see for example Davison and Hinkley, 1997). Bootstrapping allowed us to model explicitly the variation related to the incidence counts and the variation related to the leaf collection, independently of each other. This approach also allowed us to assess uncertainties in the parameter estimates without making any assumptions about the distributions of the data or the parameter values.

We created 100 000 bootstrap samples for each measurement line in each replicate in the following manner. First, we simulated the incidence counts on the measurement lines to create a distribution of incidence values. We assumed a population of 82 stems within each measurement line (based on the observed stem density) and simulated a random leaf sampling 100 000 times with all possible incidence values. We recorded the “real” incidence value each time the simulation produced the observed value. That created a distribution of “real” incidence values corresponding to our observation. Second, we sampled with replacement the analyzed leaves to generate 100 000 new samples with the original sample size. Third, the mean disease intensity of each bootstrap leaf set was multiplied by an incidence value drawn from the corresponding incidence distribution to obtain the mean intensity for each measurement line. Finally, we grouped these unconditional means over measurement lines into sets representing the five replicates. As a result, we obtained 100 000 bootstrap replicates of the entire experiment.

The one-dimensional disease gradient in equation (4) was fitted to each of the 100 000 bootstrap replicates. The disease gradient function in equation (2) was fitted to a subset of 10 000 bootstrap replicate datasets, for computational reasons. As a result, we obtained a large number of bootstrap estimates of parameters *α* and *βI*_0_ for each treatment and direction. These estimates were used to conduct statistical tests.

#### Statistical tests

Parameter differences were tested using a simple bootstrap hypothesis test (Davison and Hinkley, 1997, p. 162), where the observed difference between parameter values in different conditions is compared to a distribution of differences between those conditions in the bootstrap samples. If only a few bootstrap samples give a more extreme difference than the observed one, then the observed difference is considered significant. Significance level (*p*-value) of the observed difference is calculated by dividing the number of cases where the difference in the test statistic *t_i_* is greater than or equal to the observed difference *t_obs_* (including the observed case) by the total the number of bootstrap replicates (R) plus the observed case:

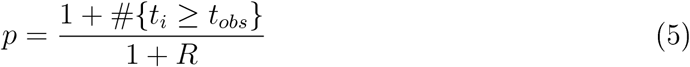

Additionally, we tested differences between conditions in the parameters *α*- and *βI*_0_ simultaneously using a two-dimensional hypothesis test based on the joint distribution of differences in *α* and *βI*_0_ (the “equidensity” test, analogous to Johansson et al., 2014). A kernel density estimate of the joint distribution was obtained to define the degree of “extremity” of a point in the two-dimensional parameter space. The point reflecting the observed difference was compared to the joint-distribution of differences between bootstrap replicates. The observed difference was considered significantly different from zero, if it was located in a sufficiently sparse area, such that less than 5% of the bootstrap estimates were located in regions with equal or lower density, analogous to being in the thin tail of a one-dimensional distribution.

We present 95% confidence intervals for the parameters derived from the distribution of bootstrap results, i.e. the limits of 2.5th and 97.5th percentile of the distribution. Additionally, differences in disease levels between treatments 1A5, 3D7 and mixed inoculation were tested at *t*_0_, *x*_0_ and *t*_1_, *x*_±1_ with the Kruskal-Wallis test and the pairwise Dunn’s posthoc comparison with the Bonferroni correction.

#### Statistics implementation

All data analysis was implemented in Python (versions 3.5.2 and 3.6.0) and the code is provided together with the data (supplement for review). Fitting was performed using lmfit-package (v. 0.9.10, Newville et al., 2014). Numerical integrations were implemented with ‘quad’ and ‘dblquad’ functions in scipy-package (v. 1.0.1, Virtanen et al., 2020). Fitting of the two-dimensional functions (Eq. 1 and 2) was performed using the high performance computing cluster Euler of ETH Zurich. Kruskal-Wallis test was conducted with ‘kruskal function in scipy-package and Dunn’s test with function ‘posthoc_dunn’ in package scikit-posthocs (v. 0.3.8, Terpilowski, 2018).

## Results

The second inoculations with strains 1A5 and 3D7, and their mixture were successful: at *t*_0_ we observed increased disease levels in the inoculation areas of all three treatments compared to controls (Fig. 2 A). At time point *t*_1_, there was a gradient of disease intensity from higher levels at *x*_±1_ to lower levels at *x*_±4_ (Fig. 2 B). Genotyping the re-isolated strains confirmed the successful spread (Appendix A). In total, 4190 plants were inspected in the course of incidence measurements, 2527 leaves were collected and analyzed using the digital image analysis, and 153 isolates were genotyped The entire dataset including raw data, bootstrap replicates, best fitting parameter estimates and weather data is available in DATADRYAD (TBA after acceptance in the journal).

### Estimates of pathogen dispersal parameters

Fitting the equation (2) to the observed disease gradients allowed us to estimate parameters *α* (dispersal parameter) and *βI*_0_ (transmission rate × initial intensity at the source). For 1A5 treatment, the estimates of *α* were very low and estimates of *βI*_0_ were very high, compared to 3D7 and mixed inoculation (Table 1). The results are biologically unrealistic (as discussed below in subsection “Estimates of disease transmission rates”) and were likely due to an insufficient disease intensity within the inoculation area and consequently a shallow gradient outside the inoculation area (Fig. 2 A and B).

**Table 1:** The best-fitting parameters from Eq. (2).

| Treatment | - direction | $\alpha$ (95% CI), cm | $\beta I_0$ (95% CI), pycnidia/leaf |
| --- | --- | --- | --- |
| 1A5 | - positive | 2.5 | 99992 |
|  | - negative | 4.9 | 3518 |
|  | - combined | 2.6 | 99975 |
| 3D7 | - positive | 15.3 (12.3 - 19.7) | 1559 (1147 - 2112) |
|  | - negative | 12.0 (9.3 - 15.5) | 2387 (1613 - 3638) |
|  | - **combined** | **13.5 (11.5 - 16.1)** | **1915 (1475 - 2430)** |
| Mixed inoculation | - positive | 23.1 (15.9 - 37.5) | 1271 (1052 - 1618) |
|  | - negative | 20.0 (14.8 - 28.4) | 1281 (1049 - 1586) |
|  | - **combined** | **21.4 (16.7 - 28.6)** | **1271 (1095 - 1468)** |

Note: Confidence intervals (CI) derived from bootstrapping, except for the implausible results of strain 1A5. Values of $\beta I_0$ were limited below 100000 in the fitting procedure.

Less successful spread of 1A5 than 3D7 was confirmed by comparing disease intensities between the treatments at *t*_0_, *x*_0_ and at *t*_1_ *x*_±1_. At *t*_0_, *x*_0_, the disease intensity was significantly lower for 1A5 than for 3D7 (Kruskal-Wallis test *H* = 10.66 *p* = 0.005, pairwise Dunn’s test *p* = 0.004). Further, at *t*_1_ *x*_±1_ the intensity in treatment 1A5 was lower than the intensity in both the 3D7 and the mixed inoculation treatments (Kruskal-Wallis *H* = 117.1 *p* = 3.4 × 10^−26^; Dunn’s test 1A5 vs 3D7 p = 6.0 × 10^−24^: Dunn’s test 1A5 vs mixed inoculation *p* =2.6 × 10^-18^); at other measurement distances the Kruskal-Wallis test detected no differences. Due to the differences in 1A5 treatment compared to others, the next steps of analysis are presented only for 3D7 and mixed inoculation treatments.

Comparison of the best-fitting parameters (Table 1) between the “positive” (North West) and the “negative” (South East) directions revealed no significant difference neither in treatment 3D7 (equidensity p-value: *p_2D_* = 0.21, one-dimensional hypothesis test Eq. (5) for parameter *α*: *p_α_* = 0.17, parameter *βI*_0_: *p_βI_0__* = 0.13, Fig. 3 A), nor in mixed inoculation treatment (*p_2D_* = 0.7% *p_α_* = 0.60, *PβI_0_* = 0.95). This similarity between directions suggests isotropic dispersal during the experiment.

**Figure 3:**
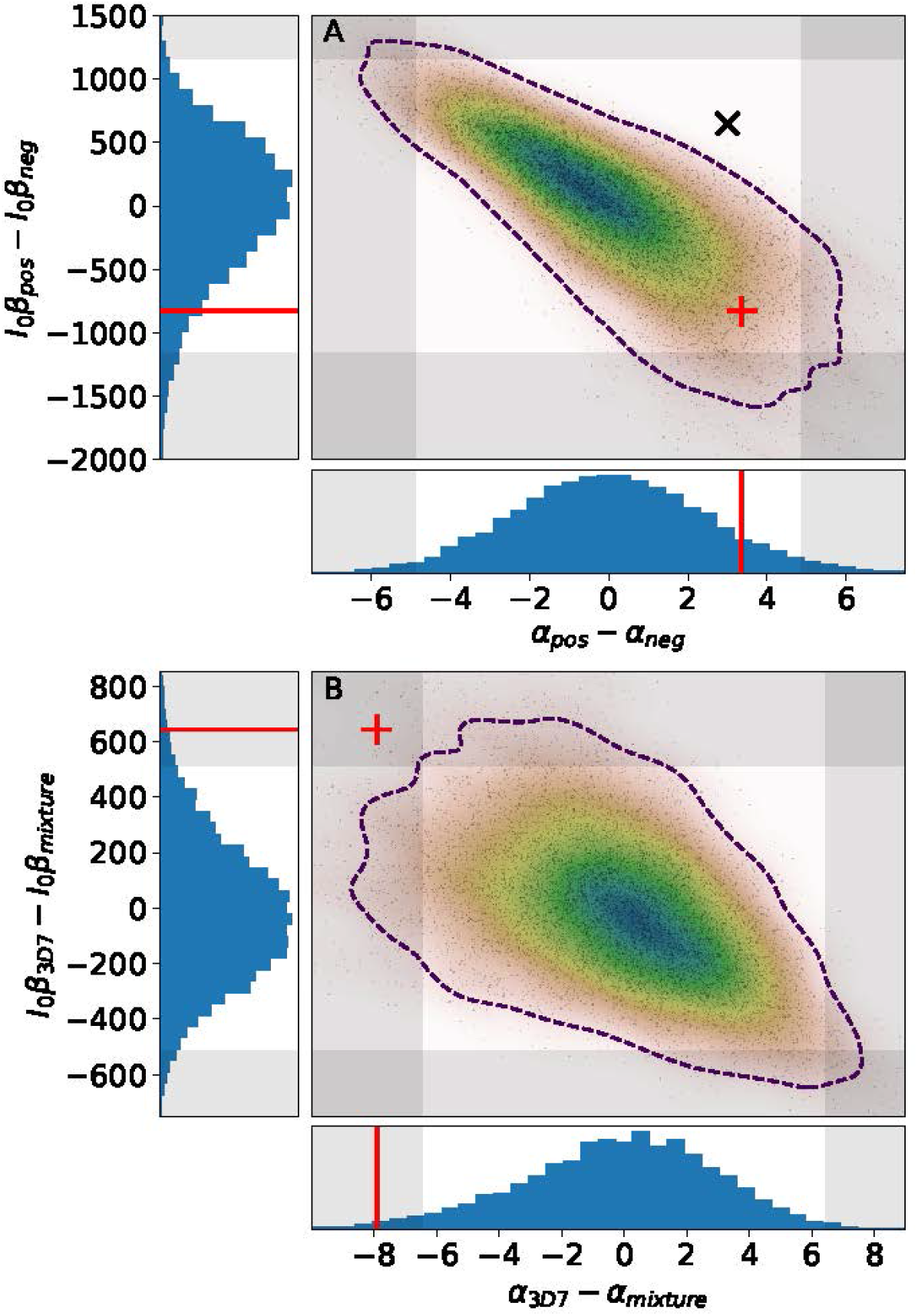
Visualization of the one- and two-dimensional bootstrap tests. (A) Comparison between two directions of dispersal of the strain 3D7. (B) Comparison between 3D7 and mixed inoculation treatments. Histograms show single parameter distributions while heat maps and black dots show joint distributions (10 000 bootstrap replicates). Observations (red line; red cross) in the 5% extreme of the distribution (shaded area; outside of the dashed line), are considered significant. The differences are significant between treatments (B) but not between directions (A). Black cross in panel (A) shows a hypothetical observation where the difference would be deemed non-significant for each parameter separately (in non-shaded area), but the joint test reveals a significant difference (outside of the dashed line).

Considering this isotropy, we estimated the parameters using the dataset that combined the two directions. We established a significant difference between 3D7 and mixed inoculation treatments (*p_2D_* = 0.014, *p_α_* = 0.020, *p_βI_0__* = 0.018, Fig. 3 B). The dispersal parameter *α* was higher in mixed inoculation while *βI*_0_ was higher in 3D7 treatment (Table 1). We thus highlight that following the mixed inoculation, the pathogen has dispersed further but caused lower disease transmission compared to 3D7 treatment.

### Estimates of disease transmission rates

In addition to estimating the pathogen dispersal parameter, we estimated the transmission rate of the disease. The fitting yielded estimates of *βI*_0_, from which we extracted the transmission rate *β* by dividing *βI*_0_ by estimates of *I*_0_. We estimated *I*_0_ = 249 pycnidia/leaf for 3D7 and *I*_0_ = 227 pycnidia/leaf for mixed inoculation measured as the mean intensity in flag leaves. Based on these estimates we calculated *β* = 7.7 (unitless) for 3D7 and *β* = 5.6 for mixed inoculation treatment.

Parameter estimates for the strain 1A5 were not realistic. It is not biologically plausible that spores of the strain 1A5 would disperse by only 3 cm, while spores of the strain 3D7 disperse some 14 cm, because pycnidiospores of the two different strains are expected to have similar physical properties. Likewise, the estimates of *βI*_0_ for strain 1A5 were unrealistically high (almost 100 000, Table 1). However, we still inferred the parameter *βI*_0_ for the strain 1A5 assuming that the physical process of spore dispersal via rain droplets is the same for the two strains. Under this additional assumption (i.e., by setting *α*_1*A*5_ = *α*_3*D*7_ = 13.5 cm), we found no difference in the spread of the strain 1A5 between the two directions (p = 0.38, parameter *βI*_0_). When we combined the data from the two directions, we estimated *βI*_0_ = 349 pycnidia/leaf and *β* = 3.0 (*I*_0_ = 118 pycnidia/leaf). The transmission rate estimates of the strain 1A5 and the mixed inoculation were lower than the estimate for the strain 3D7 (1A5 vs 3D7: *p* = 1.00 × 10^−4^, mixed inoculation vs 3D7: *p* = 0.0498).

### Genotyping as confirmatory approach

Genotyping of the 153 re-isolated strains (Appendix A) supported the conclusions drawn from the phenotypic data. (i) The inoculated pathogen strains have spread out from the inoculated area: we detected them in the measurement lines, both after inoculations with a single strain (9 isolates out of 19 were identified as strain 1A5 at *x*_±1_ for 1A5 treatment, and 45 out of 55 as 3D7 for 3D7 treatment) and after the mixed inoculation (two 1A5 isolates and 37 3D7 isolates out of the total 49 isolates sampled at *x*_±1_ versus one 1A5 isolate and eight 3D7 isolates out of total 30 isolates sampled at *x*_±3_). (ii) The proportion of identified 1A5 and 3D7 isolates was lower further away from the inoculation area in the mixed inoculation treatment. We can thus conclude that the strain 1A5 has spread less successfully than the strain 3D7: the proportion of the 3D7 isolates in the 3D7 treatment was higher than the proportion of 1A5 isolates in the 1A5 treatment and the same effect was observed in the mixed inoculation treatment.

### How good are the simplifications?

We compared the three different models (Eqs. 1, 2, 3) with regard to (i) the accuracy of the estimates and (ii) the computational time.

(i) The parameter estimates are virtually the same when using the two two-dimensional models (line destination vs. explicit destination area), which justifies our use of the slightly simplified model in Eq. (2). In contrast, the one-dimensional model resulted in substantially higher estimates of *α* (Table 2).

**Table 2:** Comparison of the parameter estimates between different functions.

| Equation | Mixed inoculation treatment B |  | 3D7 treatment |  |
| --- | --- | --- | --- | --- |
| | $\alpha$, cm | $\beta I_0$, pycn./leaf | $\alpha$, cm | $\beta I_0$, pycn./leaf |
| Eq. 1, 2D | 21.4 | 1265 | 13.5 | 1889 |
| Eq. 2, 2D | 21.4 | 1271 | 13.5 | 1915 |
| Eq. 3, 1D | 24.6 | 1131 | 15.1 | 2031 |

Note: Estimates with combined data of the two directions. “2D” and “1D” stand for two- and one-dimensional models, respectively.

Additionally, the relationship between the population spread and dispersal parameter *α* is different between one- and two-dimensional models. This difference becomes clear for example when dispersal is described based on “mean dispersal distance” which is *α* for one-dimensional exponential kernel but 2*α* for two-dimensional. For the strain 3D7, the mean dispersal distances in one- and two dimensional models would be 15 cm and 27 cm, respectively. Clearly, a one-dimensional model of dispersal should not be used for deriving dispersal distances.

(ii) Regarding the computational time, the one-dimensional model (Eq. 3) was fitted to 100 000 bootstrap datasets on a standard PC in a few hours, while fitting the two-dimensional model (Eq. 2) required a few days of computational time for only 10 000 replicates. When using the most complete model (Eq. 1) it took more than 12 hours on a PC to obtain the estimates for only one replicate (the observed data). We conclude, that taking into account spatial extent of the source by means of two-dimensional modeling pays off in the increased accuracy of the parameter estimates, even though the computational demand increases.

## Discussion

### Take home message

We measured primary, horizontal disease gradients produced by rain-splash driven dispersal and subsequent transmission via asexual spores of of *Z. tritici* in wheat canopy. We analyzed the data using a spatially-explicit mathematical model and achieved the first estimates of the dispersal kernel of the pathogen in field conditions. We report that the characteristic spatial scale of dispersal is tens of centimeters, which is consistent with previous studies in controlled conditions. Our estimation of the dispersal kernel can be used to parameterize epidemiological models that describe spatial-temporal disease dynamics within individual wheat fields.

### Measurement of the pathogen population

The use of digital image analysis allowed us to measure pycnidia numbers (representing pathogen reproduction) as a proxy for the pathogen population size. While disease severity measured as the proportion of leaf area covered by lesions of STB (PLACL) is a key quantity associated with yield loss, pathogen reproduction is more relevant for pathogen ecology and evolution. One example of this relevance is that pycnidia numbers are more powerful than PLACL for predicting the future PLACL (Karisto et al., 2018).

The dispersal kernels estimated here correspond to effective dispersal of the pathogen, not to the dispersal of all spores. A difference between those may arise from density-dependent post-dispersal mortality (Nathan et al., 2012; Klein et al., 2013). At high spore densities, such as close to the source, leaves can become saturated decreasing the infection efficiency of the spores (Karisto et al., 2019*a*). Alternatively, dispersal of the spores could be measured with spore traps placed within the canopy. However, that would leave open how many spores actually attach to healthy plants, how many of them are successful in causing lesions, and how much the established population disperses. Using healthy plants as spore traps leads to the measurement of a combination of dispersal and infection processes that is more relevant epidemiologically.

### Estimation of dispersal and transmission parameters

Dispersal parameter estimates *α*_3D7_ = 13.5cm and *α*_mix_ = 21.4cm correspond to half distances 22.7cm and 35.9 cm for 3D7 and mixed inoculation treatments, respectively (half-distance ≈ 1.7*α* in 2D Karisto et al., 2019*b*). Fitt et al. (1987) report a notably shorter range of 6-16cm for half-distances of dispersal of splash-borne fungal spores reviewing literature, including estimates for *Z. tritici* from Brennan et al. (1985), but they used a one-dimensional model of dispersal. Our one-dimensional estimates (Table 2) correspond to half-distances of 10.5 cm and 17.0 cm for 3D7 and mixed inoculation, respectively (half-distance ≈ 0.69*α* in 1D). Thus, experiments in controlled conditions appear to translate well into field but details of the models need to be taken into account for a sound interpretation of results. We computed the one-dimensional estimates solely to compare them to results of previous studies, while we consider the two-dimensional estimates to be more accurate.

While the horizontal dispersal measurements made in previous experimental studies were useful, they are difficult to extrapolate to real field conditions for two reasons. First reason is that the modeling approaches did not consider the spatial extent of the source area as we did here but assumed a unique point source (Karisto et al., 2019 *b*). Second reason is that plant canopy, acting as a barrier that limits the effective dispersal distance, was not always included in the design (e.g. Brennan et al., 1985).

The estimated transmission rates were *β*_1*A*5_ = 3.0, *β*_3*D*7_ = 7.7, and *β*_mix_ = 5.6. The intermediate transmission of the mixed inoculation likely resulted from the contrasting reproductive capacities of the two strains. Strain 1A5 is known to produce fewer and smaller pycnidia than strain 3D7 on cultivar Runal in greenhouse (Stewart et al., 2018), which is further confirmed in field conditions by our results.

STB development depends on weather conditions (Henze et al., 2007). Also, the disease levels within the source and along the disease gradient were measured only on upper leaf layers, but dispersal occurred likely to, and possibly also from, the lower leaf layers as well, which were not included in our estimates of transmission rates. For all these reasons, the relative differences between transmission rate estimates in our experiment are informative, while their comparison with transmission rates measured in other studies is of limited value.

### Differences in dispersal

According to our analysis, the mixture of the two strains has spread further than the single strain 3D7. The difference may arise from various sources. Density-dependent effects may have flattened the disease gradient in the mixed inoculation treatment, for example if the mixed pathogen population is more sensitive to saturation at high population densities close to the inoculated area due to cross-suppression between the two pathogen strains. In this case, the beginning of the gradient, where the population density is highest, would experience stronger saturation after mixed inoculation compared to single strain inoculations. That would result in lower overall transmission but also flatter gradient (i.e., longer dispersal), as the tail of the gradient would be relatively stronger after the mixed inoculation compared to the single strain inoculations due to relaxed cross-suppression. Interestingly, these expected patterns matched with our observations in the present study.

Also, sexual reproduction in the mixed inoculation treatment could result in a shallower gradient due to wind-dispersed ascospores which are expected to have a much higher dispersal distance. The average latent period following infections by ascospores was shown to be longer than the latent period associated with asexual pycnidiospores (Morais et al., 2015), but the distribution of ascospore latent periods might still be wide enough to contribute to our observed gradient (Suffert and Thompson, 2018). Ultimate causes of the difference remain unknown.

### Sampling distances

The measurement lines were at closest 20±5 cm from the edge of the inoculated area. Measuring the disease intensity closer to the source and even inside the source might have improved the estimates, as the differences between the gradients are more pronounced close to the source. However, closer to source the reliability of measurements may suffer from saturation and also from dispersal via direct contact (Fitt et al., 1989).

We measured the disease also inside the inoculation area, but those data points were excluded from fitting, because our aim was to analyze only the newly spread infection and capture the primary disease gradients. Increase of the disease intensity in the source area from *t*_0_ to *t*_1_ was not only due to secondary infections but likely also from latent infections that can become symptomatic after a long time (Karisto et al., 2019*a*). An additional reason for excluding data at *x*_0_ is that saturation effects are expected to be strongest there.

### Spatially-explicit modeling

The spatially-explicit modeling that accounted for the spatial extent of the source allowed us to parameterize the actual dispersal kernel, while overcoming the practical issues related to a point source. We were able to create sufficiently strong, measurable disease gradients in two out of three treatments. Further extension of the source might have created a sufficiently strong gradient for strain 1A5 too. Overall, using an extended source provides a major improvement for the purpose of measuring dispersal over small scales, where using a point-source is practically impossible. The spatially-explicit modeling also allowed to estimate transmission rates β in a biologically meaningful manner. Benefits of the two-dimensional modeling compared to the one-dimensional fitting are hence undeniable.

### Statistical advances

The bootstrapping methods allowed us to test for differences in our parameter estimates and the associated uncertainties in a robust manner without making any assumptions about underlying distributions. Computationally intensive bootstrapping with a large number of replicates (100 000) is possible using modern computing resources. Non-parametric bootstrapping is a useful alternative to standard parametric tests, which are often used in biology even when their assumptions are violated. Moreover, we adopted a two-dimensional hypotheses test based on kernel density estimate of the bootstrap parameter distribution. This allowed us conduct joint testing for differences in two parameters simultaneously, which is likely to be more robust than the combination of two one-dimensional tests for each of the parameters separately. This is demonstrated in the example where a combination of 1D-tests would fail to reject the null-hypothesis (Fig. 3A). The provided source code and raw data will facilitate the application of these methods to other contexts.

### Implications for epidemiology and control

According to our estimates, the median dispersal distance of new infection from a focal source was around 30 cm. However, the limit of 95% of new infections would extend up to one meter (Karisto et al., 2019 *b*, >Eq. A3). In the second dispersal step, the infection will not only spread further outwards, but the secondary spread within the area covered by the first step would likely cause significant damage (Shaw and Royle, 1993) and this is when a clearly visible focus of the disease can be seen (Zadoks and van den Bosch, 1994). Thus, when one observes a visible focus in the canopy, the spread has likely already happened at least one step further, considering the latent infections. This should be taken into account when aiming for spatially targeted control of crop diseases in the context of precision agriculture, e.g. based on aerial detection and quantification of symptoms (Yu et al., 2018; Anderegg et al., 2019). The safety margins around a disease focus depend on the expected dispersal distance of the pathogen, which highlights the importance of precise characterization of dispersal of plant pathogens in field conditions.

### Outlook

Natural populations of *Z. tritici* are extremely diverse both phenotypically and genetically (Karisto et al., 2018; Hartmann et al., 2017). Hence, it would be interesting to conduct similar measurements in a number of other *Z. tritici* strains under various conditions in the field in order to better understand the diversity of dispersal and transmission processes. The methodology developed here can also be applied to other plant pathosystems. We hope that our study with the available raw data and source code of the analyses will lead to further measurements of dispersal of plant pathogens.

## Acknowledgements

PK and AM gratefully acknowledge financial support from the Swiss National Science Foundation through the Ambizione grant PZOOP3_161453. Genetic Diversity Centre at ETH Zurich helped with the genetic analysis. PK and AM would like to thank Andreas Hund and Hansueli Zellweger for the maintenance of the field experiment, Bruce McDonald for helpful discussions and Marcello Zala for support in preparing inoculations and other laboratory work. AM is grateful to Jonathan Levine for his advice on designing the experiment. PK is grateful to Fanny Hartmann for help in designing PCR primers. FS would like to thank Christophe Montagnier, Sandrine Gélisse and Nicolas Lecutier for maintenance of the similar field experiment prepared at the facilities of INRAE Bioger in Thiverval-Grignon, France.

## Appendix A: Genotyping the strains with PCR

### Primer design

We designed four primer pairs targeted at each of the two strains. The primers were first aimed to be fully specific to the target isolates 1A5 and 3D7 within the set of four commonly used lab strains 1A5, 3D7 1E4 and 3D1 (ST99CH_1E4, ST99CH_3D1, Zhan et al., 2002) and second, as specific to the target strain as possible in the field. Specificity here means that the primers designed for 1A5 should produce an amplicon in PCR only with 1A5 genome and not with other strains. Strain specific primers would allow for a convenient detection of the focal sub-population after the experiment as in a mark-recapture experiment.

To design the primers, we used presence-absence data of predicted genes from Hartmann and Croll (2017). We chose the target regions that were present in the target strain (either 1A5 or 3D7) and absent in the other three isolates (1E4, 3D1, and either 3D7 or 1A5). From those potential targets, we selected ten regions that were least frequently present in the 27 Swiss isolates analyzed by Hartmann and Croll (2017). After selecting the target regions, we designed four primer pairs that would be suitable for high throughput qPCR in similar conditions: amplicon length 100-150 bp, melting temperature around 60 °C. The four primer pairs were designed to amplify regions in different chromosomes of the target strain to minimize the possibility of finding all of them in a single strain in the field. Details of the designed primers are given in Table A1.

### Validation of primer specificity

First validation of the primers was done with qPCR among the four strains 3D7, 1A5, 3D1 and 1E4 (Tables A2 and A3, Figures A1 and A2). Successful amplification of the target DNA and no amplification on non-target DNA suggested that each of the eight primer pairs was specific to their target strain among the four strains, indicating successful primer design based on the genomes.

**Table A1:**
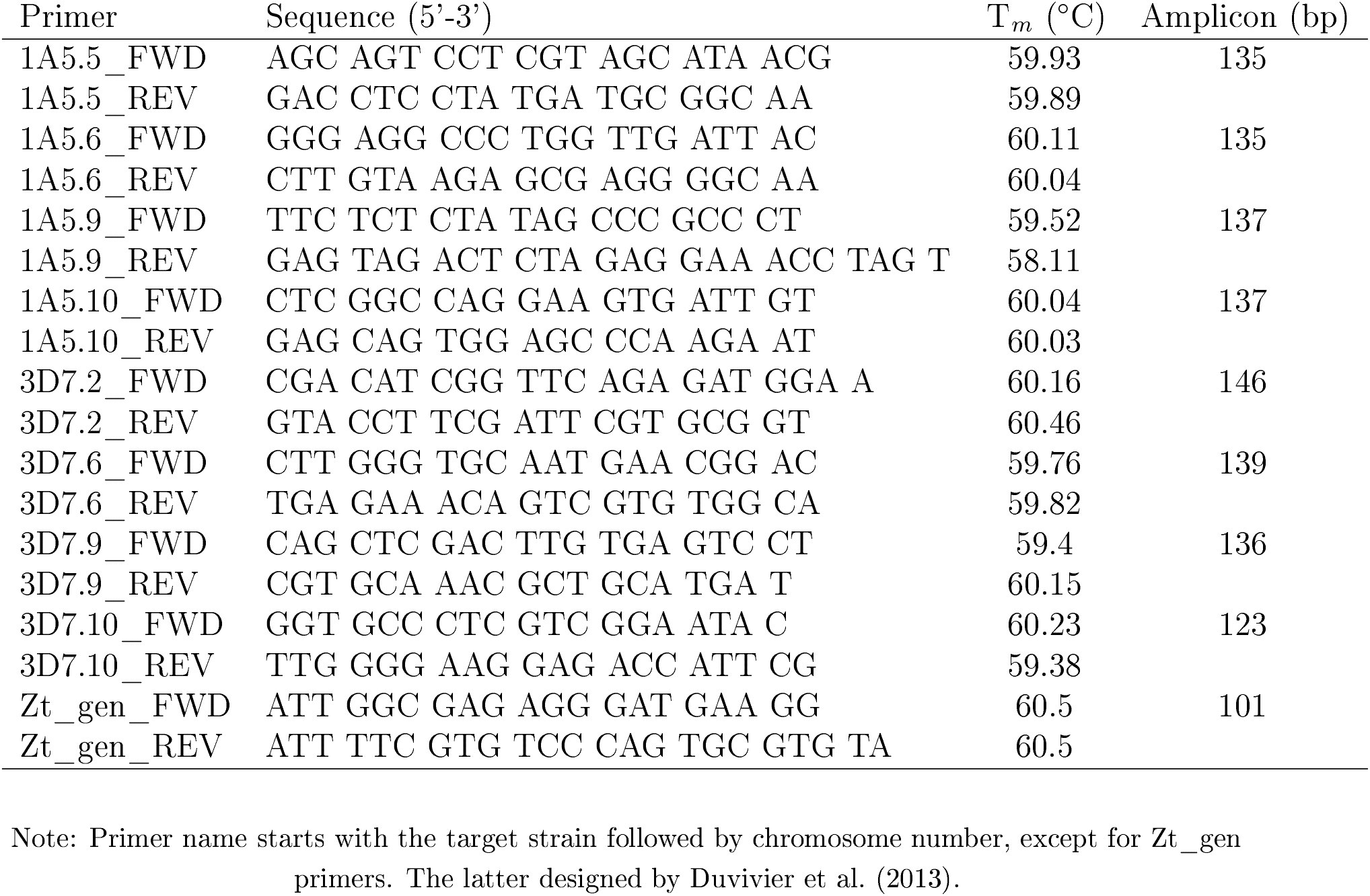
Primers

**Table A2:**
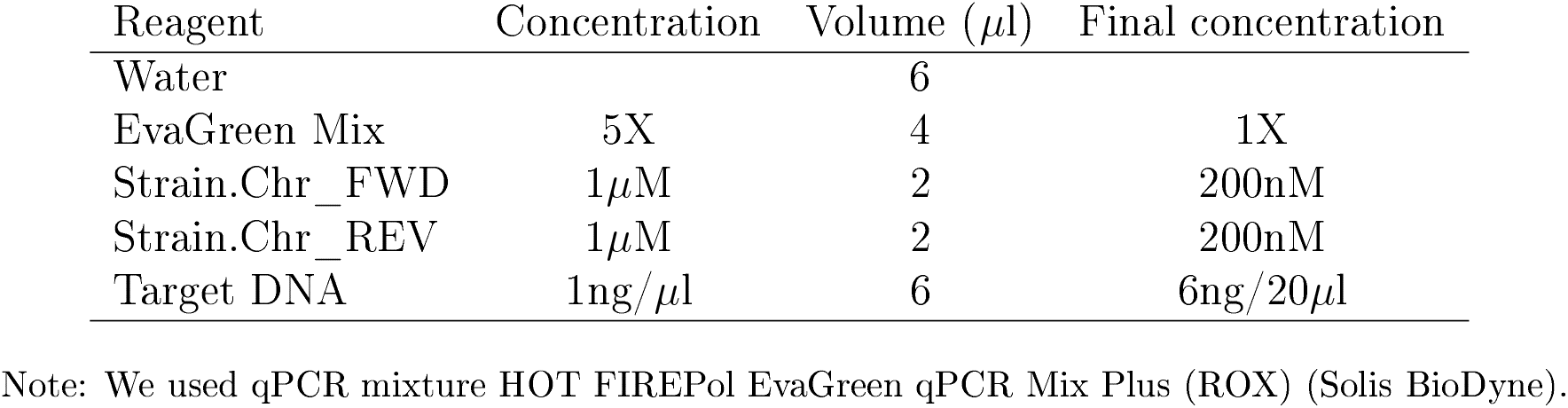
qPCR reaction mix, 20*μ*l

Primers’ specificity was then validated in a natural population using multiplex-PCR (Table A4, Table A5) combining each strain-specific primer pair with a primer pair that is specific to *Z. tritici* generally (Zt_gen primers, Duvivier et al., 2013). Zt_gen provided a positive control for success of the PCR. Primers were tested against 37 natural strains isolated from the control plots of the experiment. Reaction with primers 1A5.9 did not work reliably, indicated by the lack of Zt_gen amplicon. Numbers of false positives for other primer pairs were 4, 8, 20 for 1A5.5, 1A5.6, 1A5.10 and 13, 12, 7, 6 for 3D7.2. 3D7.6, 3D7.9 and 3D7.10 respectively (Figures A3, A4, A5, A6, A7, A8, A9, A10). Importantly, none of the false positives of 1A5.5 and 1A5.6 overlapped with each other, hence using combined data of those two gave no false positives. The six false positives of 3D7.10 were amplified with all the other 3D7-primers and none of the 1A5 primers.

**Table A3:**
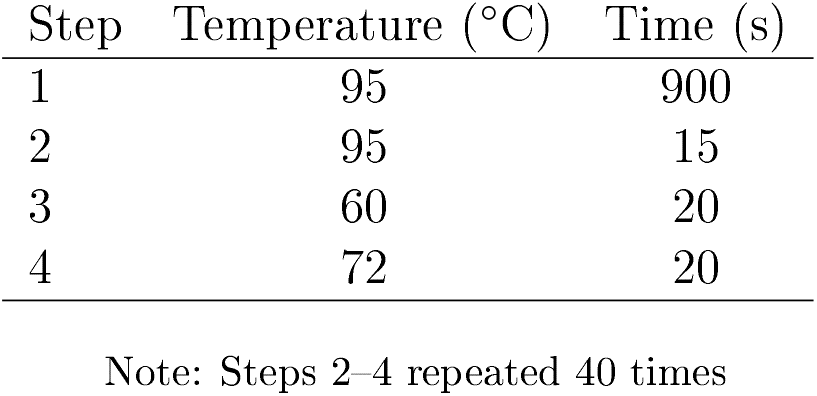
qPCR reaction cycles

| Step | Temperature (°C) | Time (s) |
| --- | --- | --- |
| 1 | 95 | 900 |
| 2 | 95 | 15 |
| 3 | 60 | 20 |
| 4 | 72 | 20 |

Note: Steps 2–4 repeated 40 times

**Table A4:** PCR reaction mix, 20*μ*l

| Reagent | Concentration | Volume (μl) | Final concentration |
| --- | --- | --- | --- |
| Water |  | 6.34 |  |
| KAPA Buffer | 2X | 10 | 1X |
| Zt_gen_FWD | 10μM | 0.5 | 250nM |
| Zt_gen_REV | 10μM | 0.5 | 250nM |
| Strain.Chr_FWD | 5μM | 1 | 250nM |
| Strain.Chr_REV | 5μM | 1 | 250nM |
| KAPA3G Polymerase | 2.5U/μl | 0.16 | 2U/100μl |
| Spore solution | 10<sup>4</sup>-10<sup>6</sup>sp/ml | 2 |  |

Note KAPA3G Plant PCR Kit (Kapa Biosystems).

**Table A5:** PCR reaction cycles

| Step | Temperature (°C) | Time (s) |
| --- | --- | --- |
| 1 | 96 | 180 |
| 2 | 95 | 20 |
| 3 | 60 | 15 |
| 4 | 72 | 15 |
| 5 | 72 | 30 |

Note: Steps 2–4 repeated 35 times.

**Figure A1:**
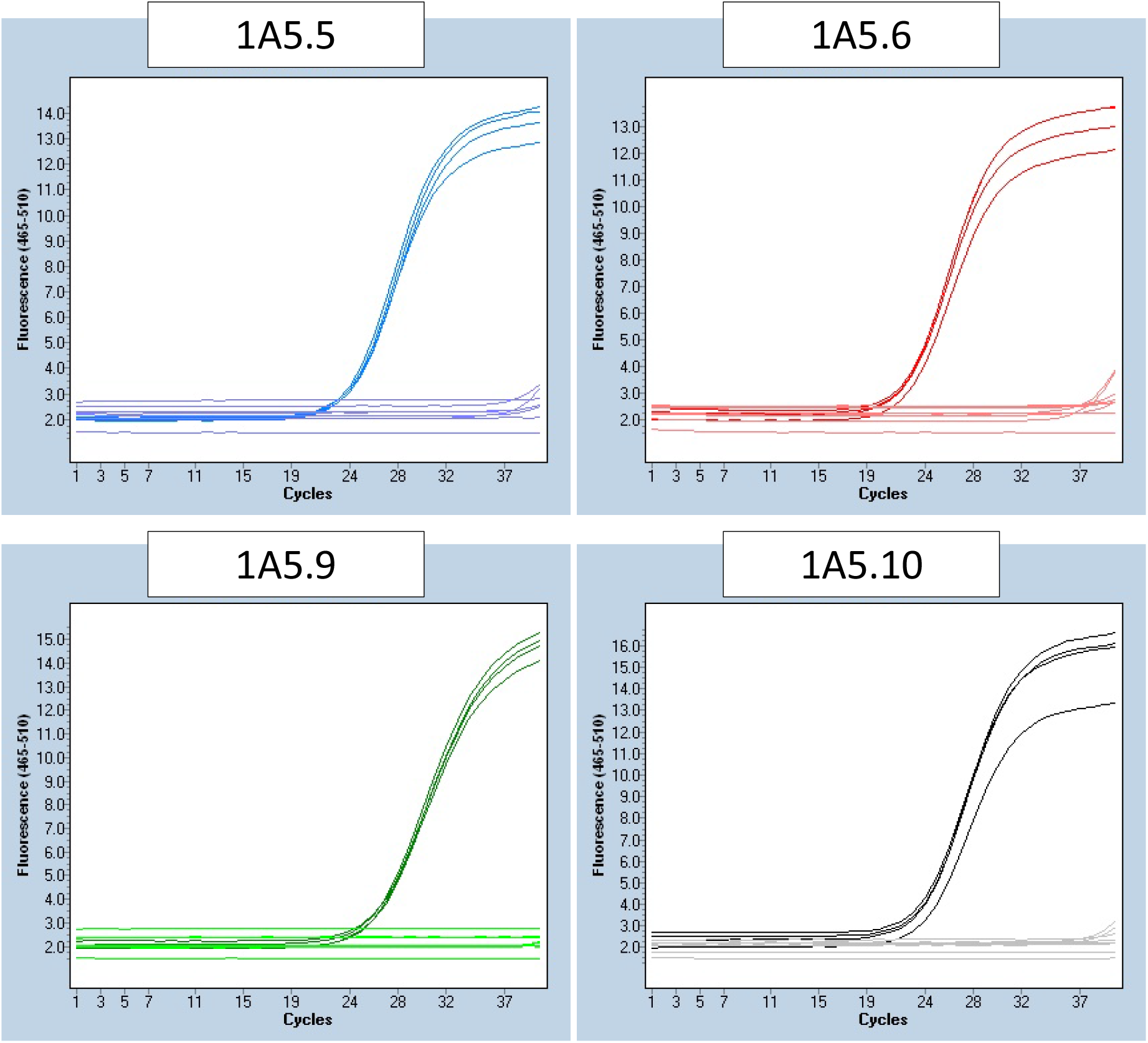
Amplification plots of the 1A5 targeting primers in qPCR. The amplified curves represent four replicates of 1A5, while the lower curves represent two replicates of each of 1E4, 3D1, 3D7 and water.

Thus, it is possible that they were the actual strain 3D7 either left on the field from previous years of field experiments or it was a spill-over from the current treatments.

**Figure A2:**
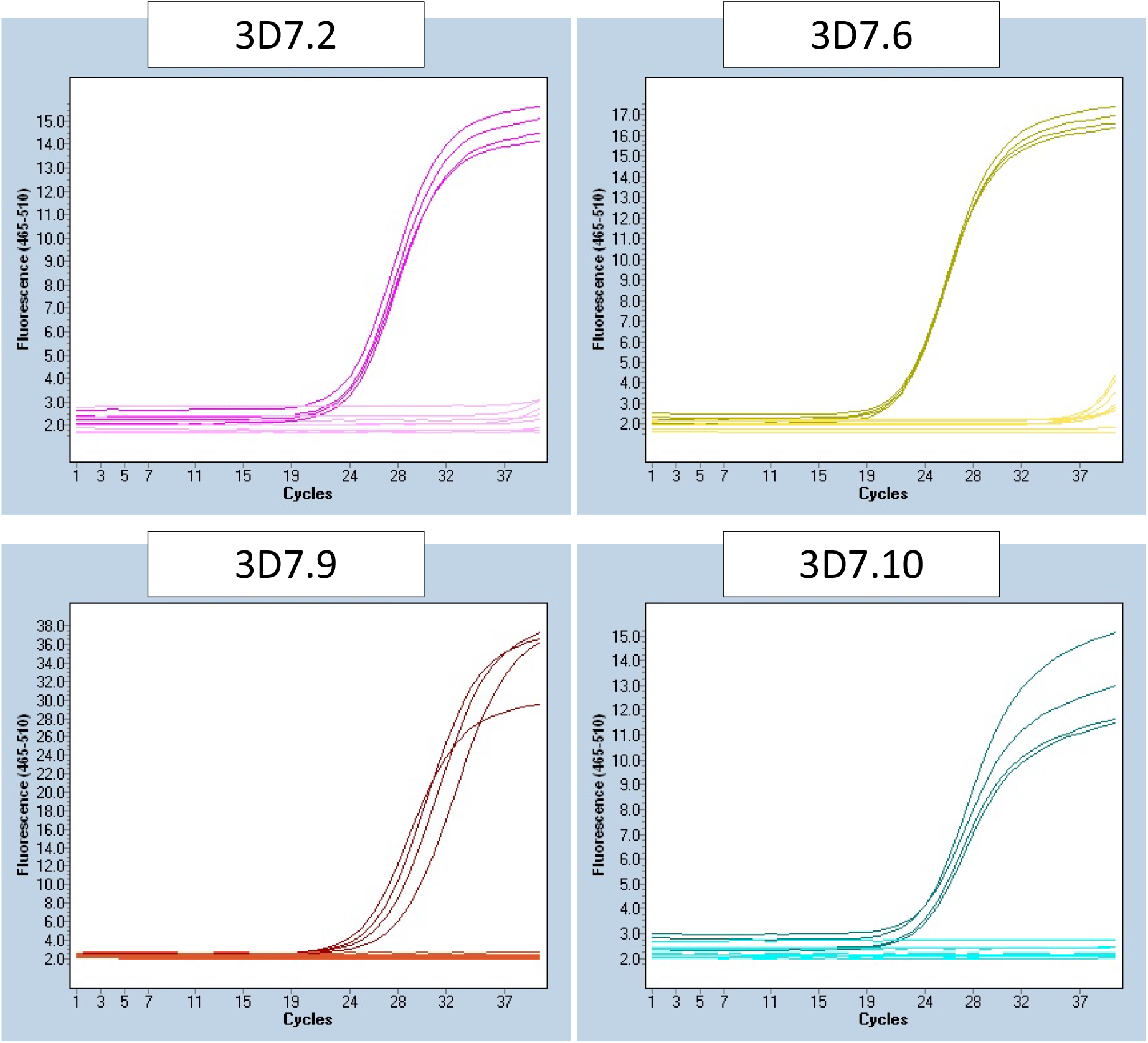
Amplification plots of the 3D7 targeting primers in qPCR. The amplified curves represent four replicates of 3D7, while the lower curves represent two replicates of each of 1A5, 1E4, 3D1 and water.

**Figure A3:**
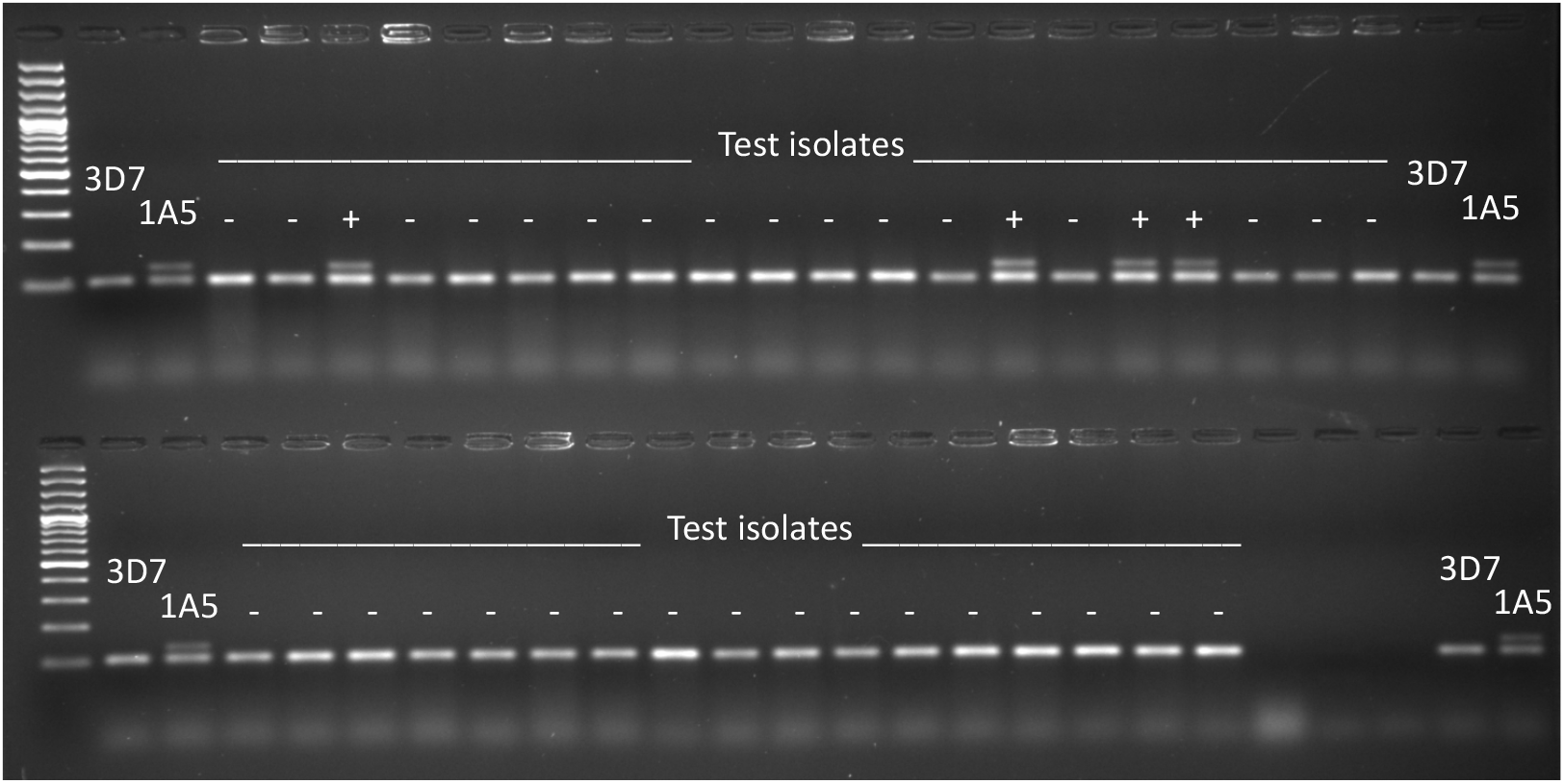
Amplicons from Zt_gen (shorter) and 1A5.5 (longer). Isolate 1A5 as positive control and 3D7 as negative control. Plus and minus indicate successful reaction (Zt_gen amplicon) and presence or absence of target amplicon, respectively.

**Figure A4:**
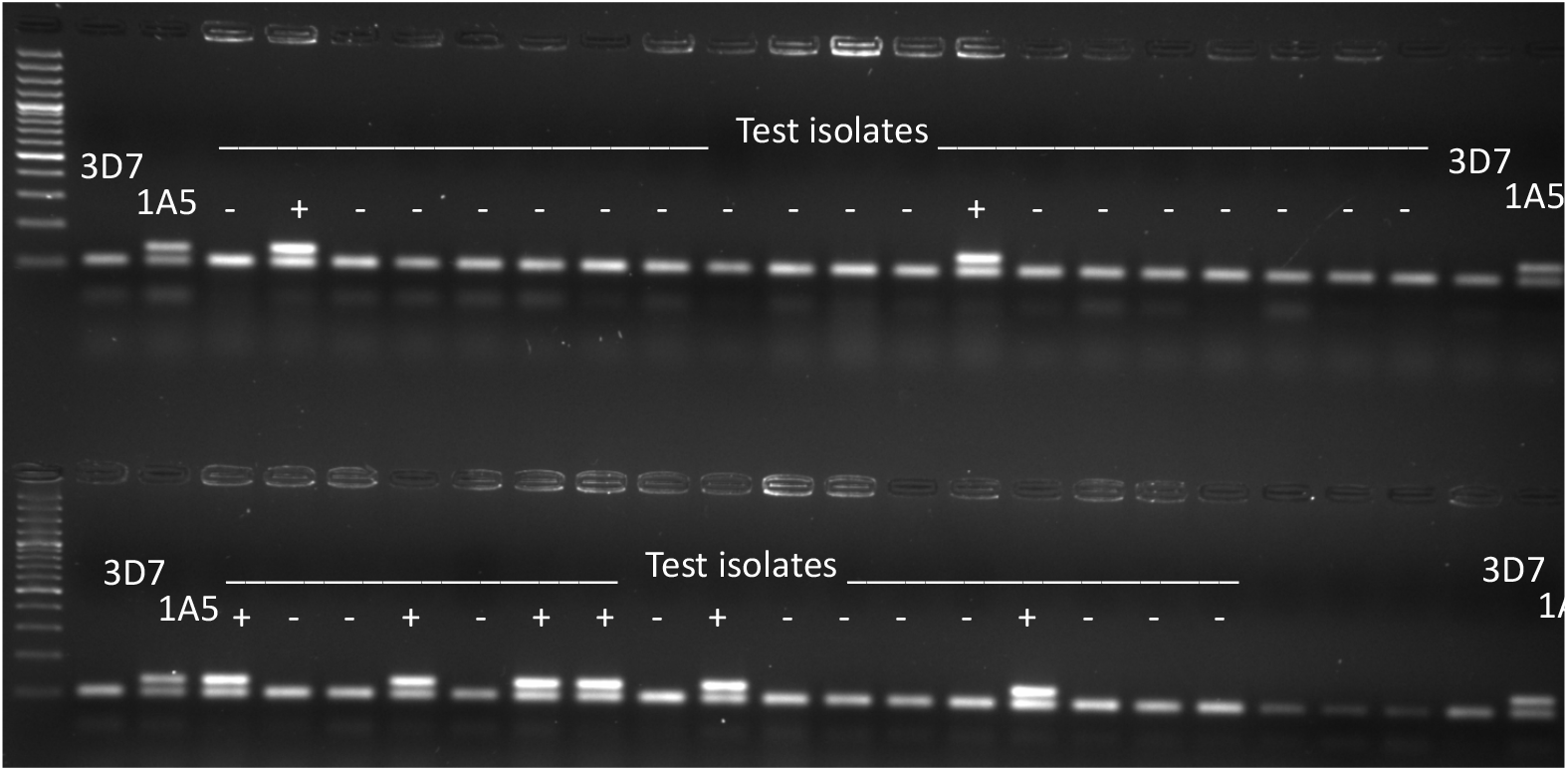
Amplicons from Zt_gen (shorter) and 1A5.6 (longer). Isolate 1A5 as positive control and 3D7 as negative control. Plus and minus indicate successful reaction (Zt_gen amplicon) and presence or absence of target amplicon, respectively.

**Figure A5:**
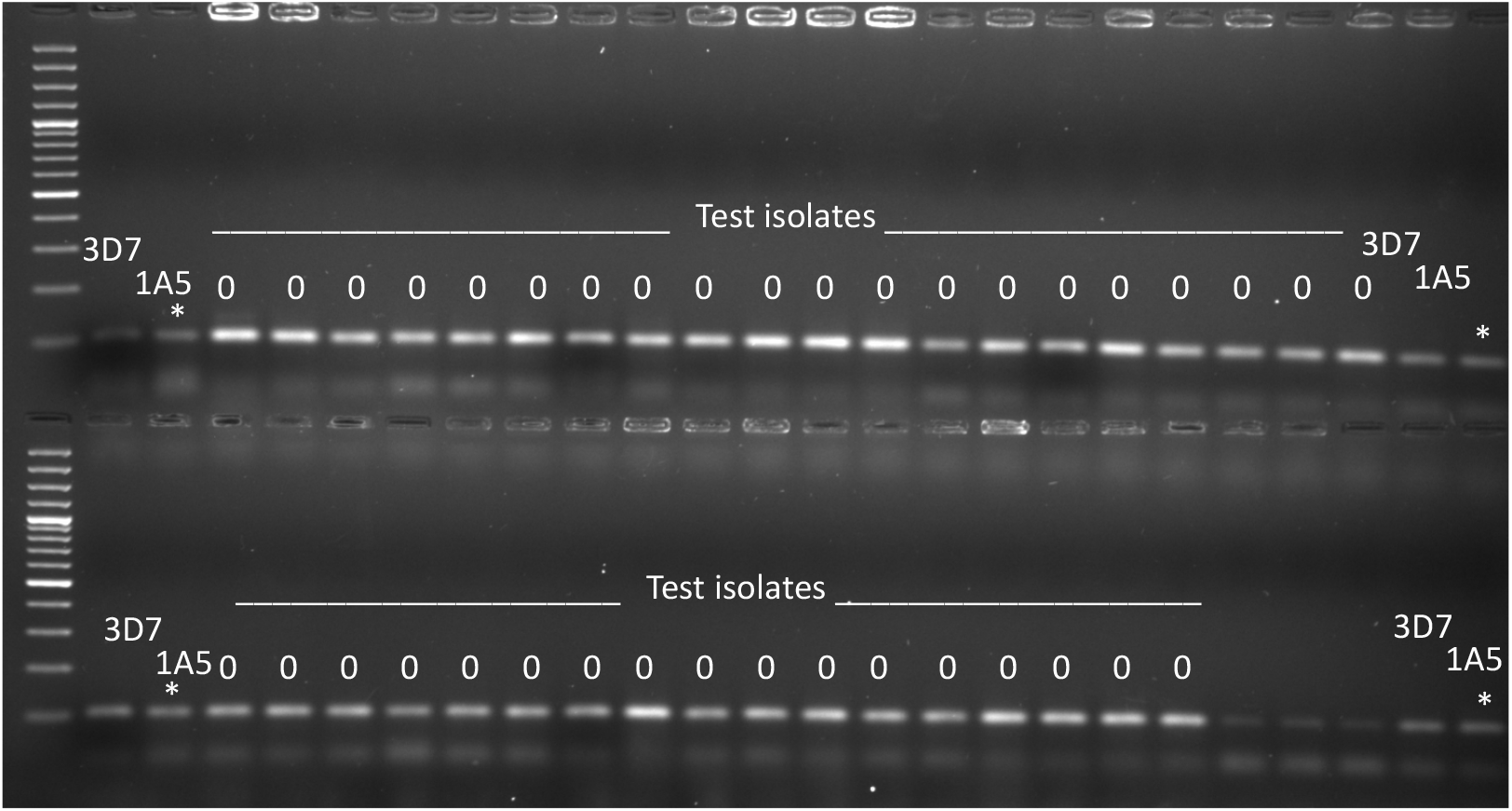
Amplicons from Zt_gen (shorter) and 1A5.9 (longer, not present). Isolate 1A5 as positive control and 3D7 as negative control. Zeros indicate that no conclusions were drawn from the reactions, as the positive controls were not amplified (*).

**Figure A6:**
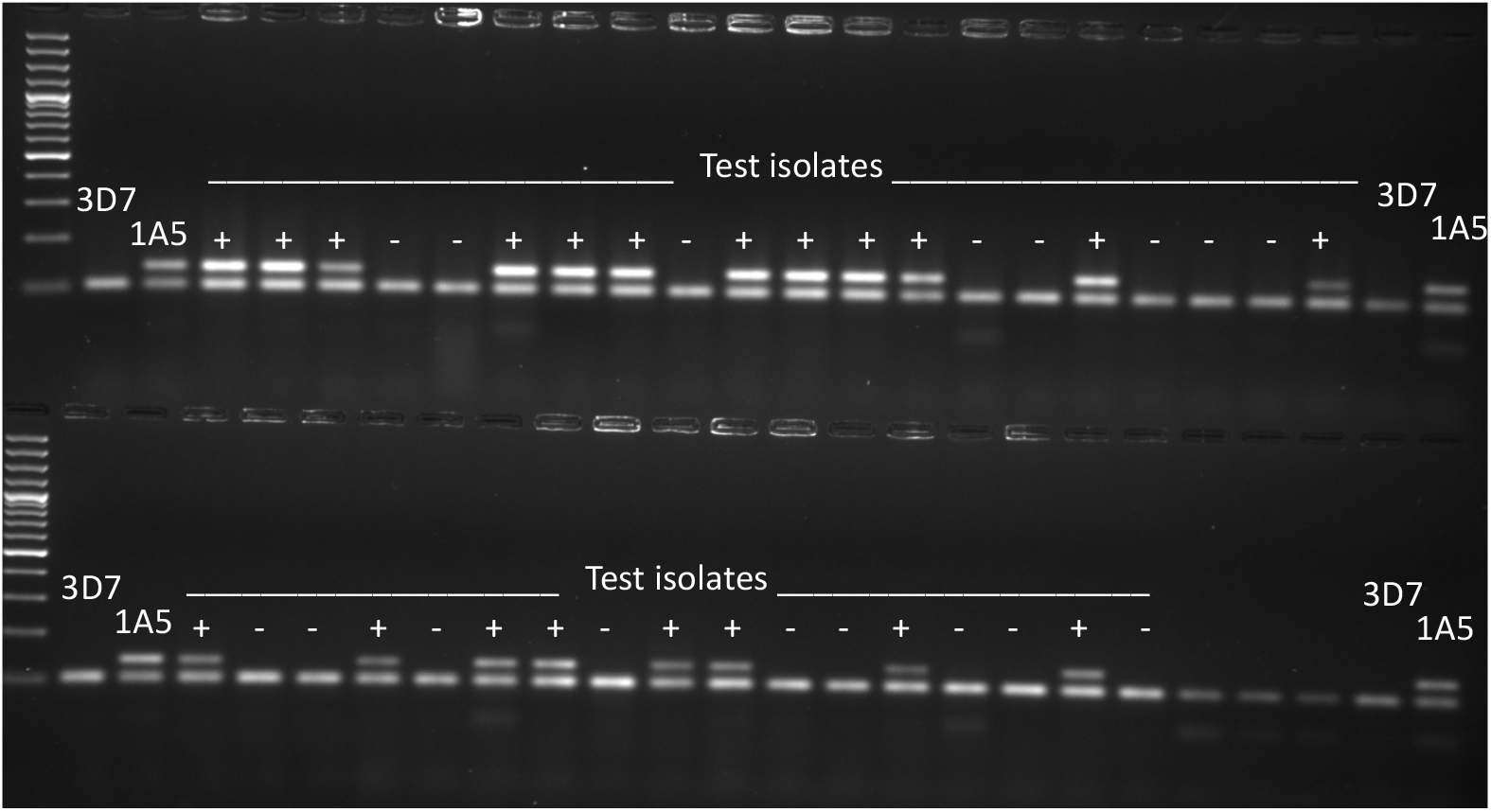
Amplicons from Zt_gen (shorter) and 1A5.10 (longer). Isolate 1A5 as positive control and 3D7 as negative control. Plus and minus indicate successful reaction (Zt_gen amplicon) and presence or absence of target amplicon, respectively.

**Figure A7:**
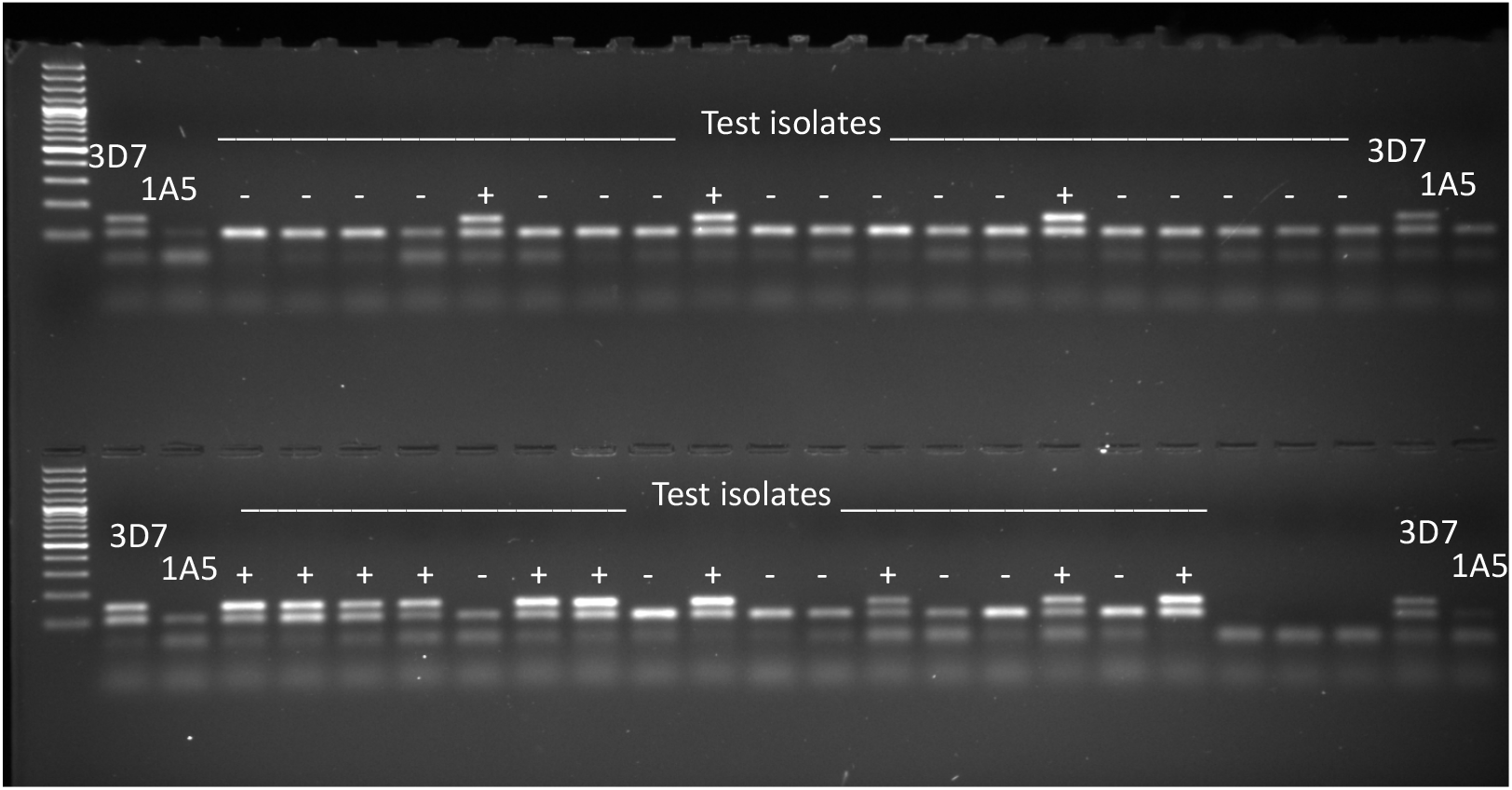
Amplicons from Zt_gen (shorter) and 3D7.2 (longer). Isolate 3D7 as positive control and 1A5 as negative control. Plus and minus indicate successful reaction (Zt_gen amplicon) and presence or absence of target amplicon, respectively.

**Figure A8:**
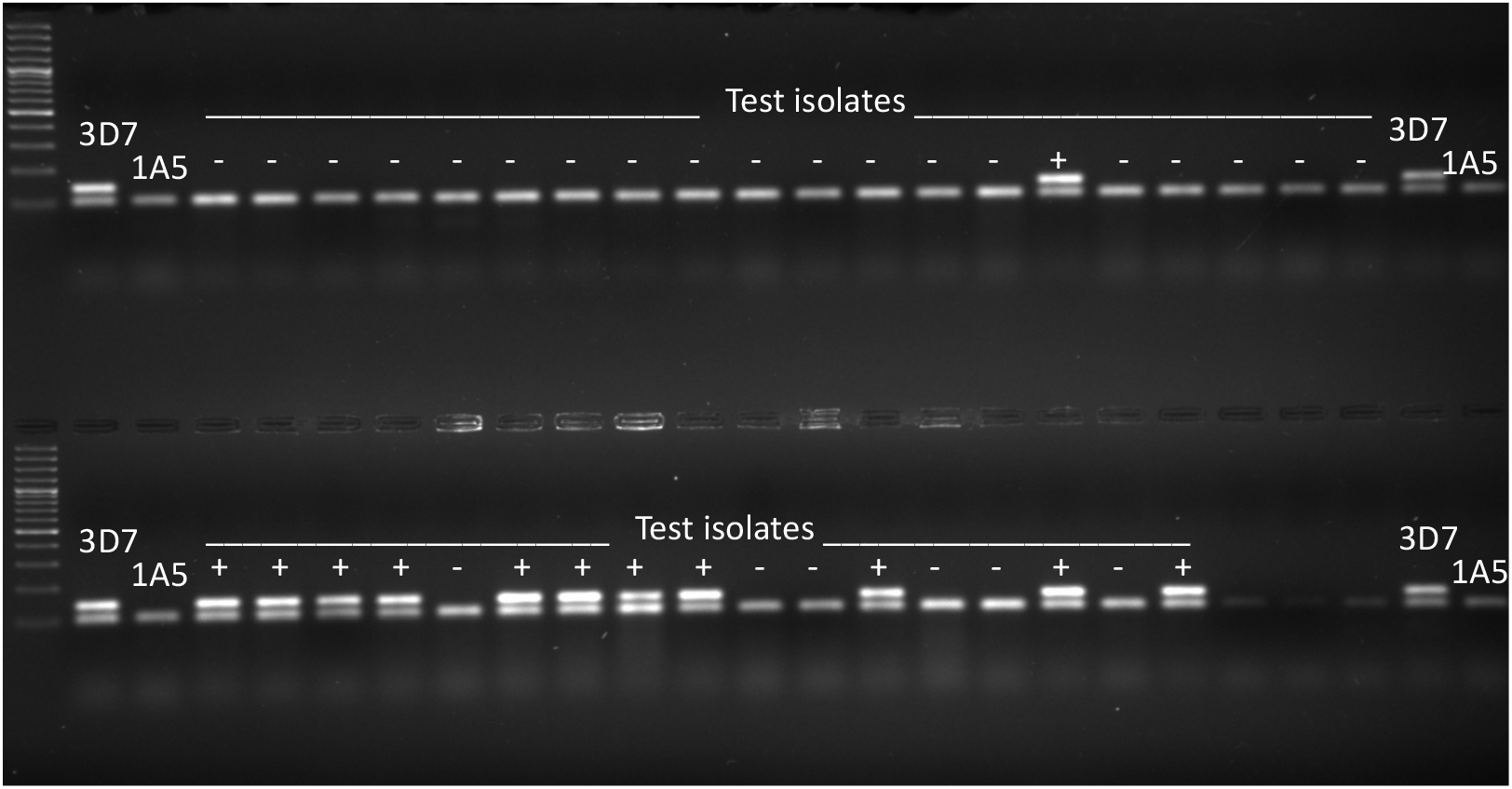
Amplicons from Zt_gen (shorter) and 3D7.6 (longer). Isolate 3D7 as positive control and 1A5 as negative control. Plus and minus indicate successful reaction (Zt_gen amplicon) and presence or absence of target amplicon, respectively.

**Figure A9:**
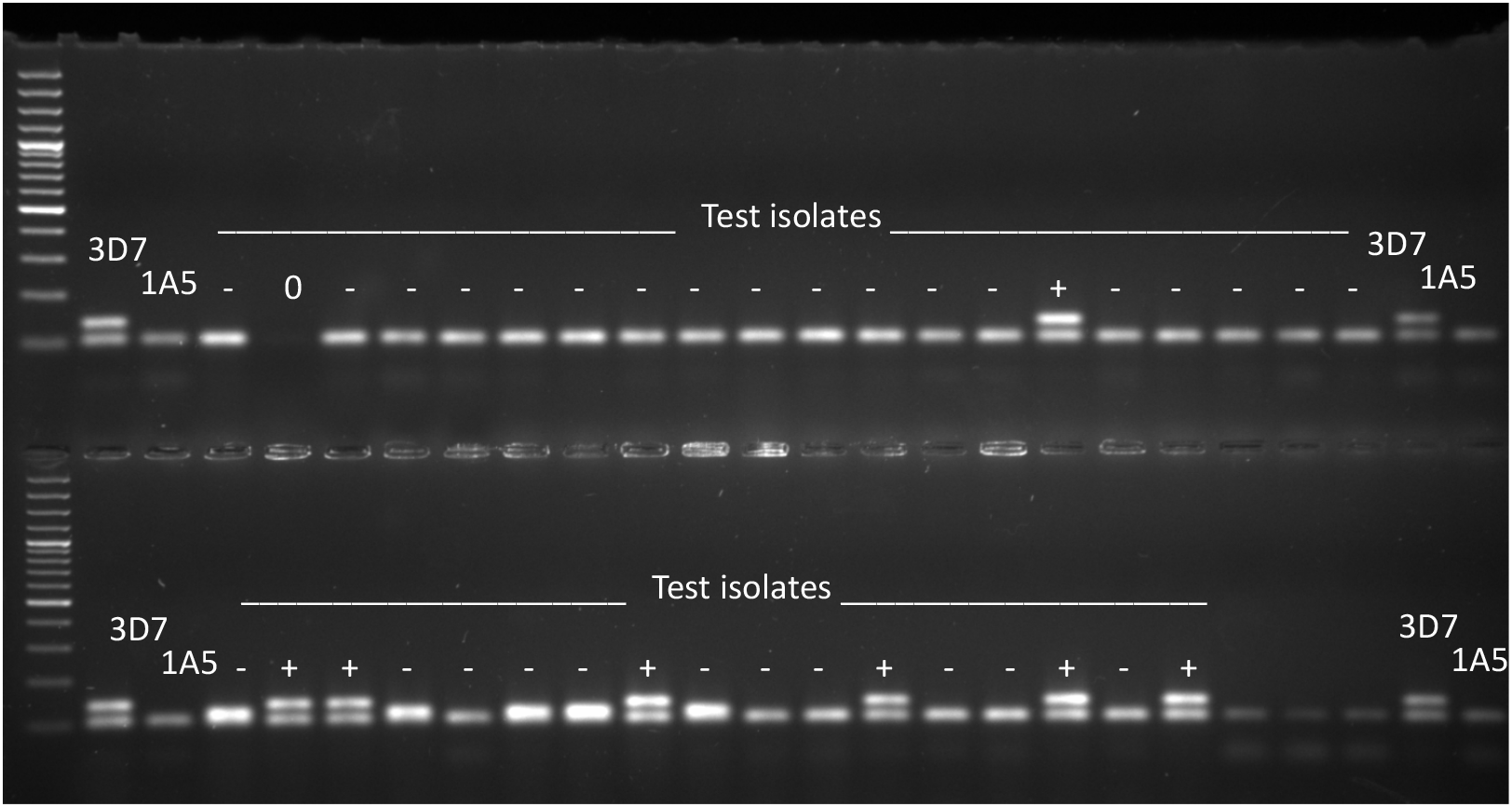
Amplicons from Zt_gen (shorter) and 3D7.9 (longer). Isolate 3D7 as positive control and 1A5 as negative control. Plus and minus indicate successful reaction (Zt_gen amplicon) and presence or absence of target amplicon, respectively.

**Figure A10:**
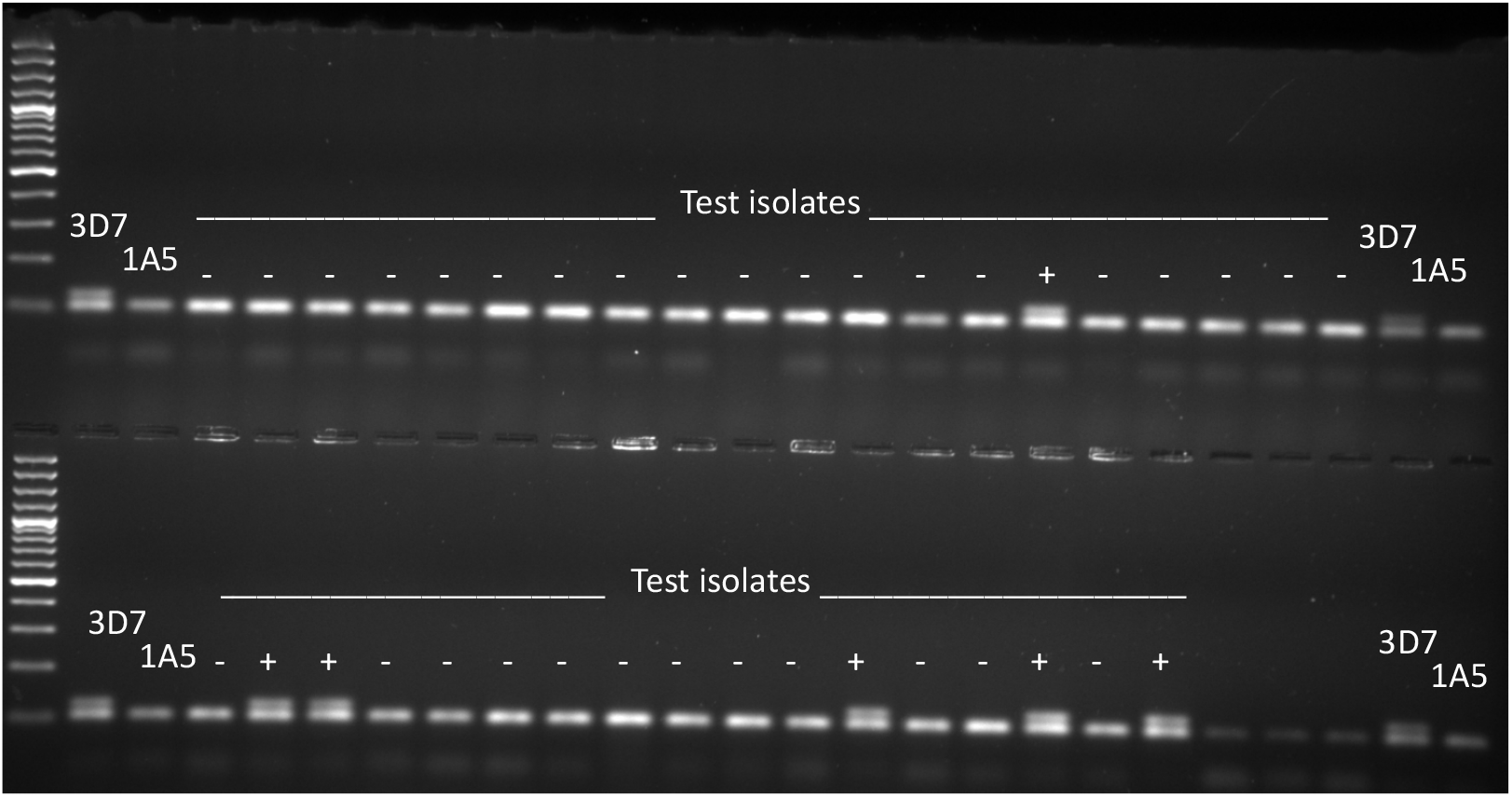
Amplicons from Zt_gen (shorter) and 3D7.10 (longer). Isolate 3D7 as positive control and 1A5 as negative control. Plus and minus indicate successful reaction (Zt _gen amplicon) and presence or absence of target amplicon, respectively.

### Genotyping of the re-isolated strains

After validation, the primers 1A5.5, 1A5.6, 3D7.9 and 3D7.10 were chosen for the genotyping the strains isolated from the leaf samples. The criterion for “detection” of one of the inoculated strains was when both of the two primer pairs targeting either 3D7 or 1A5 showed amplification. On control plots, 6/37 strains tested were detected as 3D7 (16% false positives) while 0/37 strains were detected as 1A5 (0% false positives). In a plot of treatment 1A5 (replicate 1), 9/19 strains (47%) at the first measurement lines *x*_±1_ were detected as 1A5 (Fig. A11). In contrast, in a plot of treatment 3D7 (replicate 1), 45/55 strains (82%) at *x*_±1_ were detected as 3D7 (Figs. A12, A13). Thus, frequency of 3D7 was higher than 1A5 outside the inoculation area, as implied by the disease gradients (Fig. 2B). In the plot of mixed inoculation treatment (replicate 1), at *x*_±1_ 2/49 were 1A5 and 37/49 were 3D7, while at the third measurement lines *x*_±3_ 1/30 was 1A5 and 8/30 were 3D7 (Figs. A14, A15 for 1A5, and Figs. A16, A17 for 3D7). The proportion of the target strains decreased with distance. Lower proportion of 1A5 is likely a result of two-fold effect of weaker transmission: first, the strain produced fewer pycnidia in the inoculation area (mixed inoculation treatment, replicate 1, at *x*_0_ *t*_0_: 1A5 4/15, 3D7 10/15, Figs. A11, A12, A13) and second, those pycnidia multiplied themselves with lower success.

**Figure A11:**
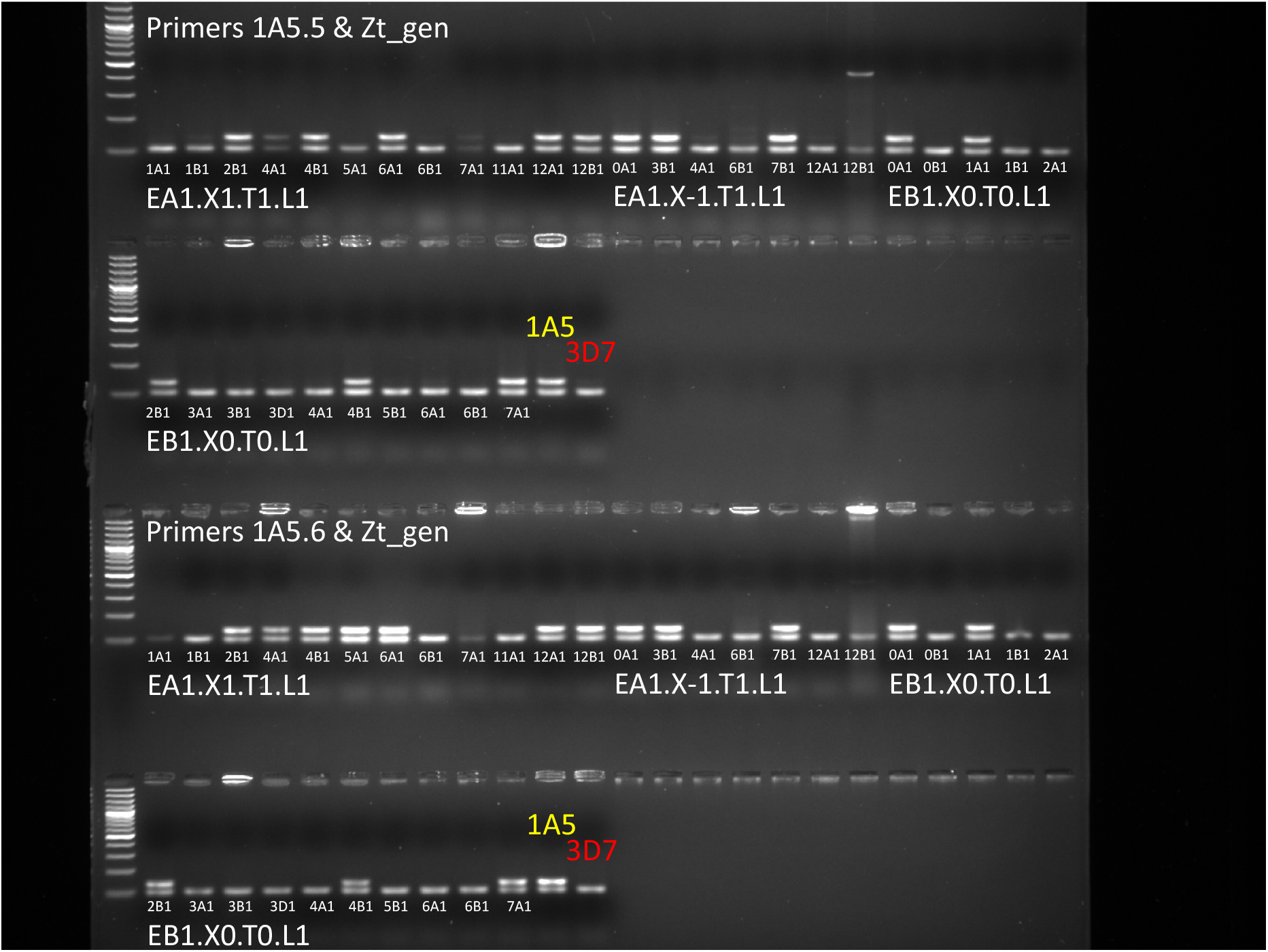
Amplicons from Zt_gen (shorter), 1A5.5 (longer, upper rows) and 1A5.6 (longer, lower rows). Isolate 1A5 as positive control and 3D7 as negative control. First part of isolate labels consist of location (E = Eschikon), treatment (e.g. A), and replicate (e.g. 1); second part contains measurement line (x1 for *x*_+1_); third, time point (T1 for *t*_1_); fourth, leaf layer (L1 = Flag); and finally the isolate itself (e.g. 1A1: leaf 1, area A, isolate 1).

**Figure A12:**
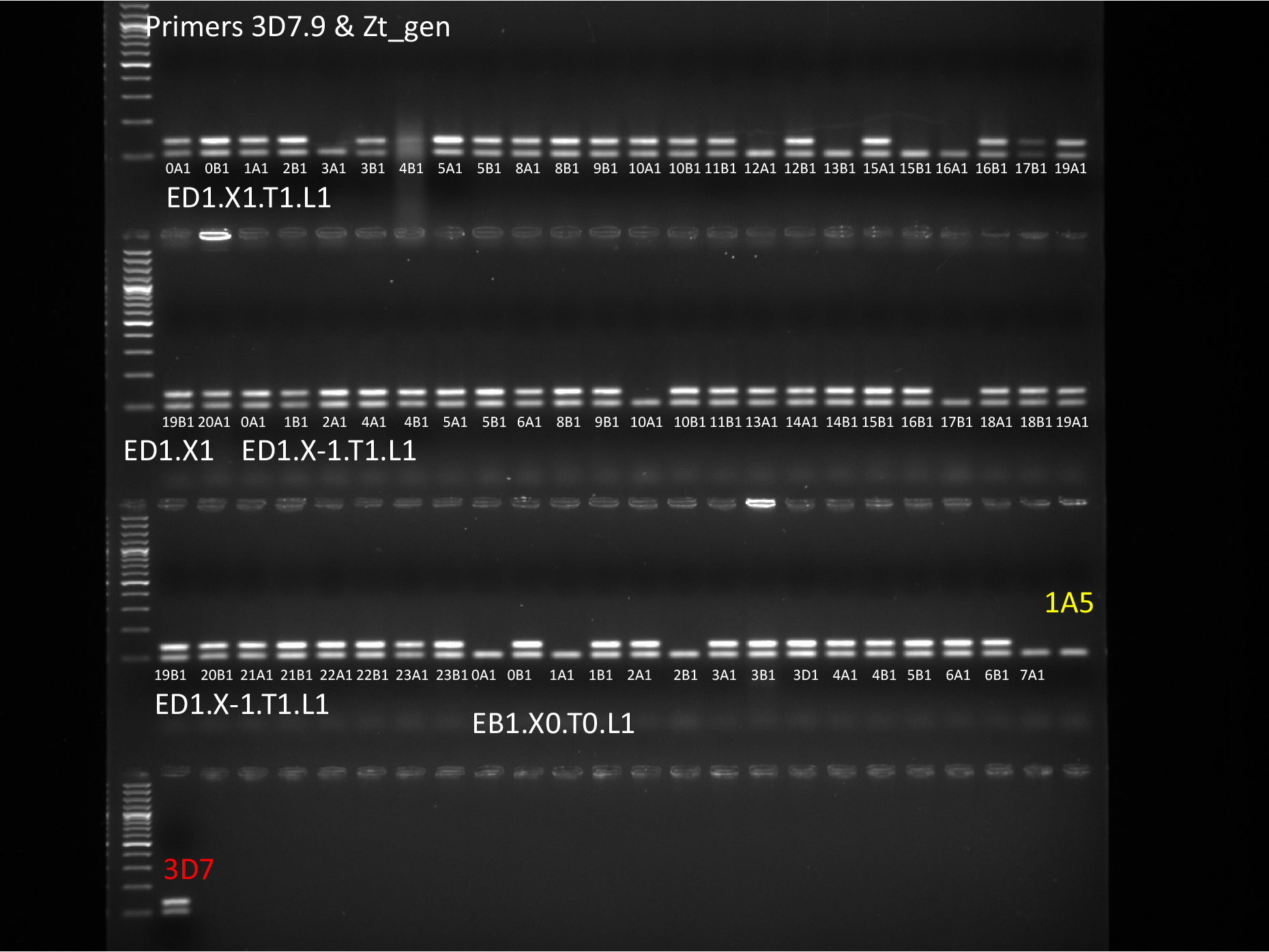
Amplicons from Zt_gen (shorter), 3D7.9 (longer). Isolate 3D7 as positive control and 1A5 as negative control. See Fig. A11 for label decoding.

**Figure A13:**
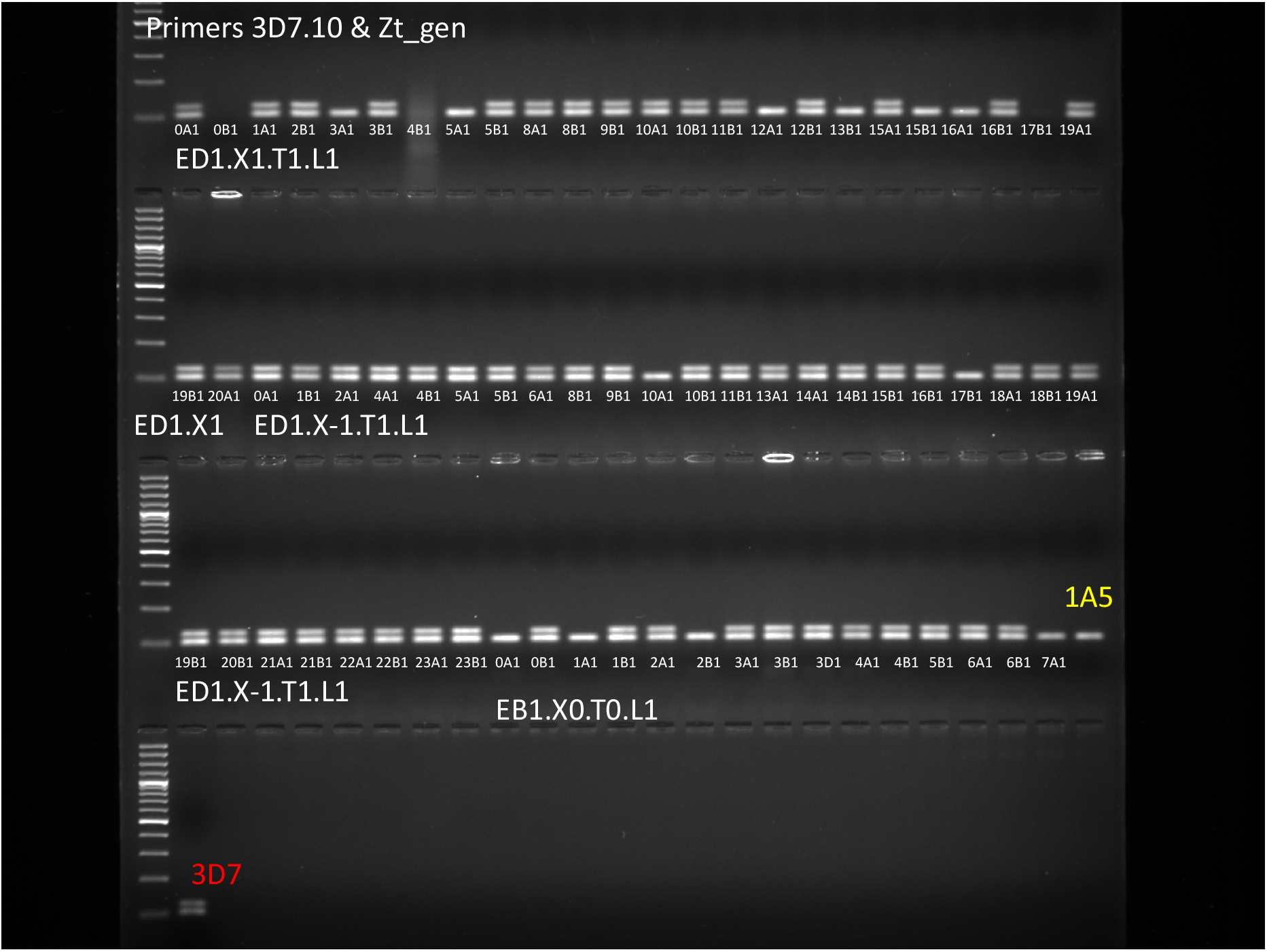
Amplicons from Zt_gen (shorter), 3D7.10 (longer). Isolate 3D7 as positive control and 1A5 as negative control. See Fig. A11 for label decoding.

**Figure A14:**
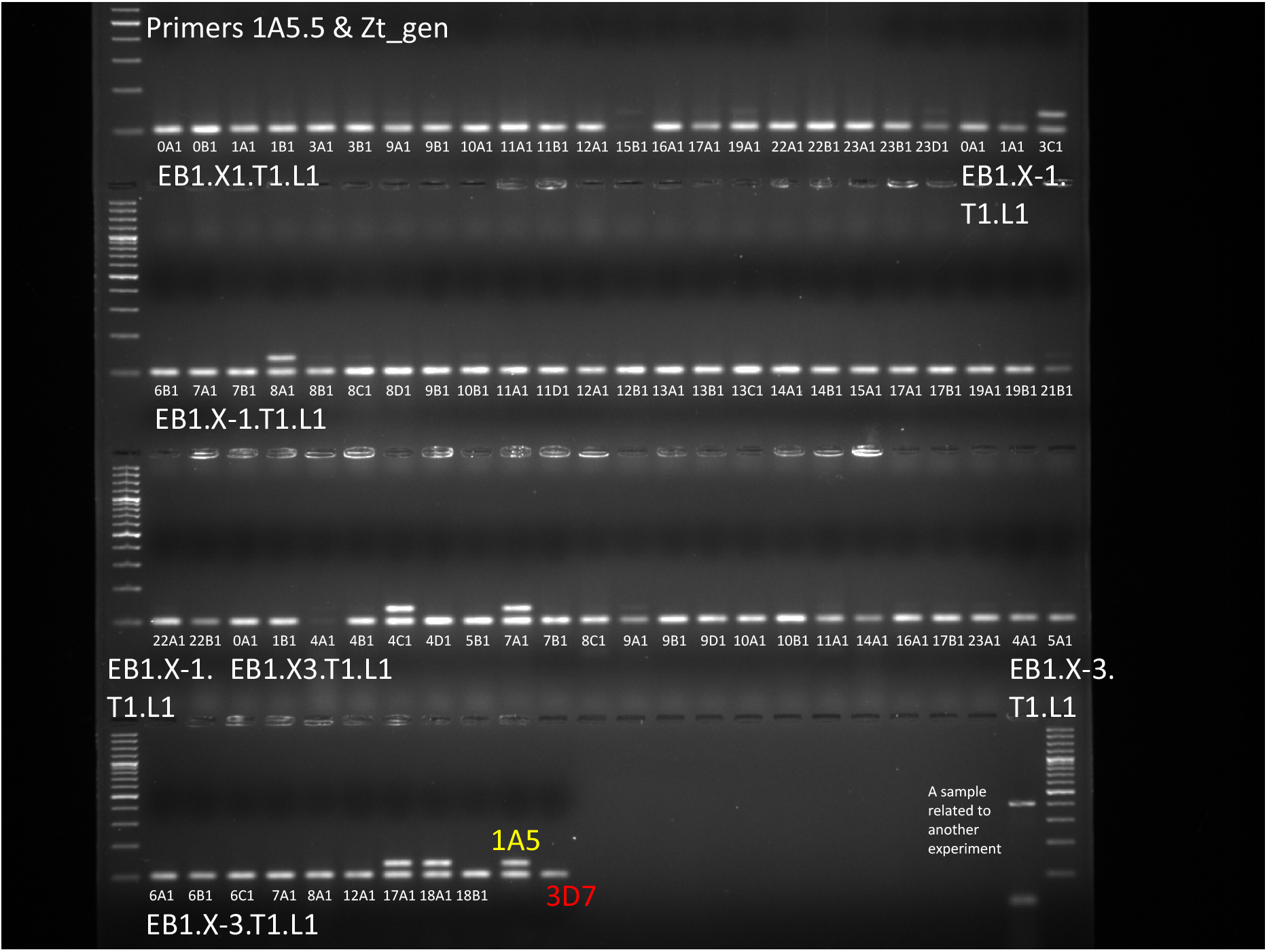
Amplicons from Zt_gen (shorter), 1A5.5 (longer). Isolate 1A5 as positive control and 3D7 as negative control. See Fig. A11 for label decoding.

**Figure A15:**
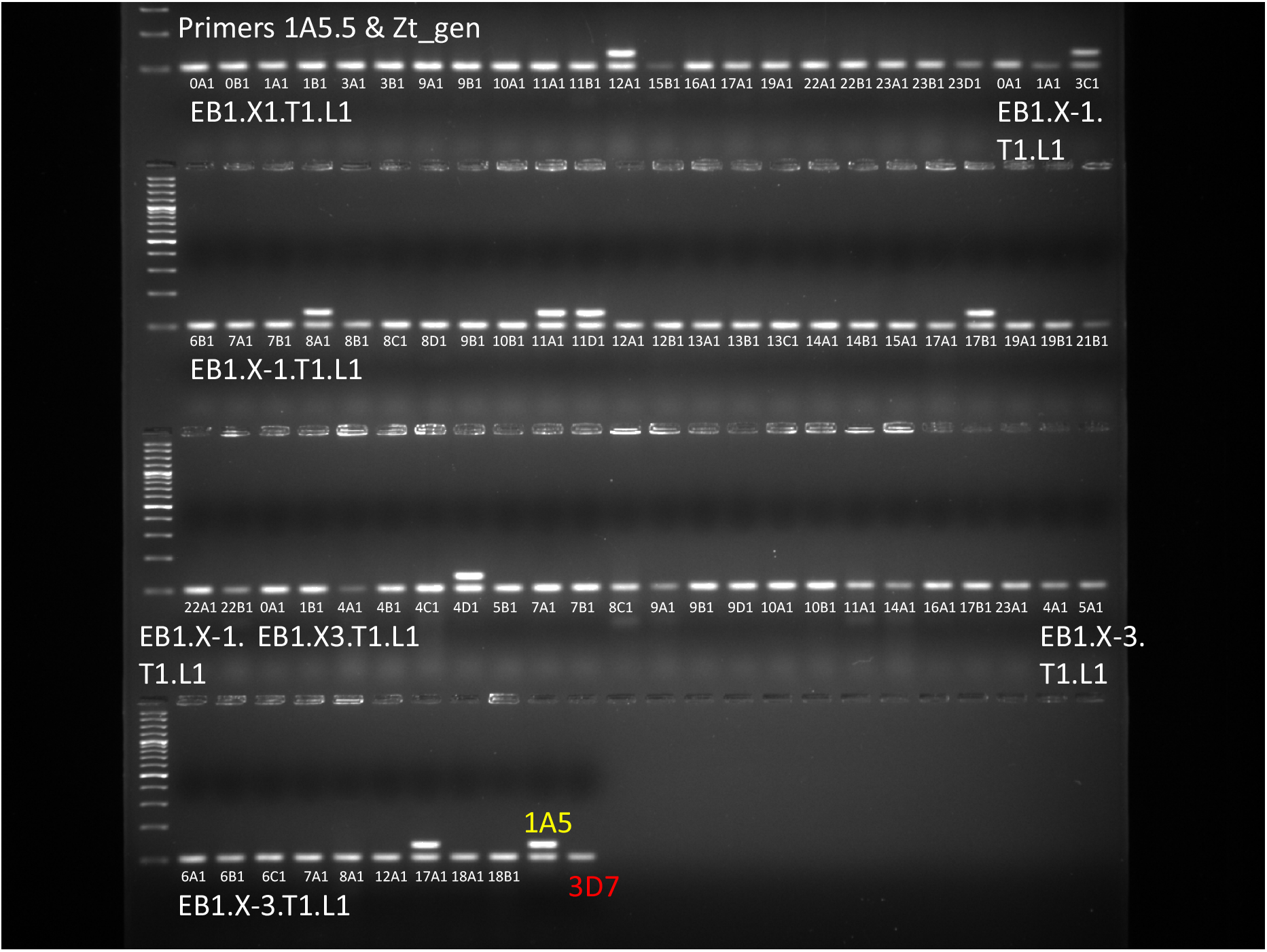
Amplicons from Zt_gen (shorter), 1A5.6 (longer). Isolate 1A5 as positive control and 3D7 as negative control. See Fig. A11 for label decoding.

**Figure A16:**
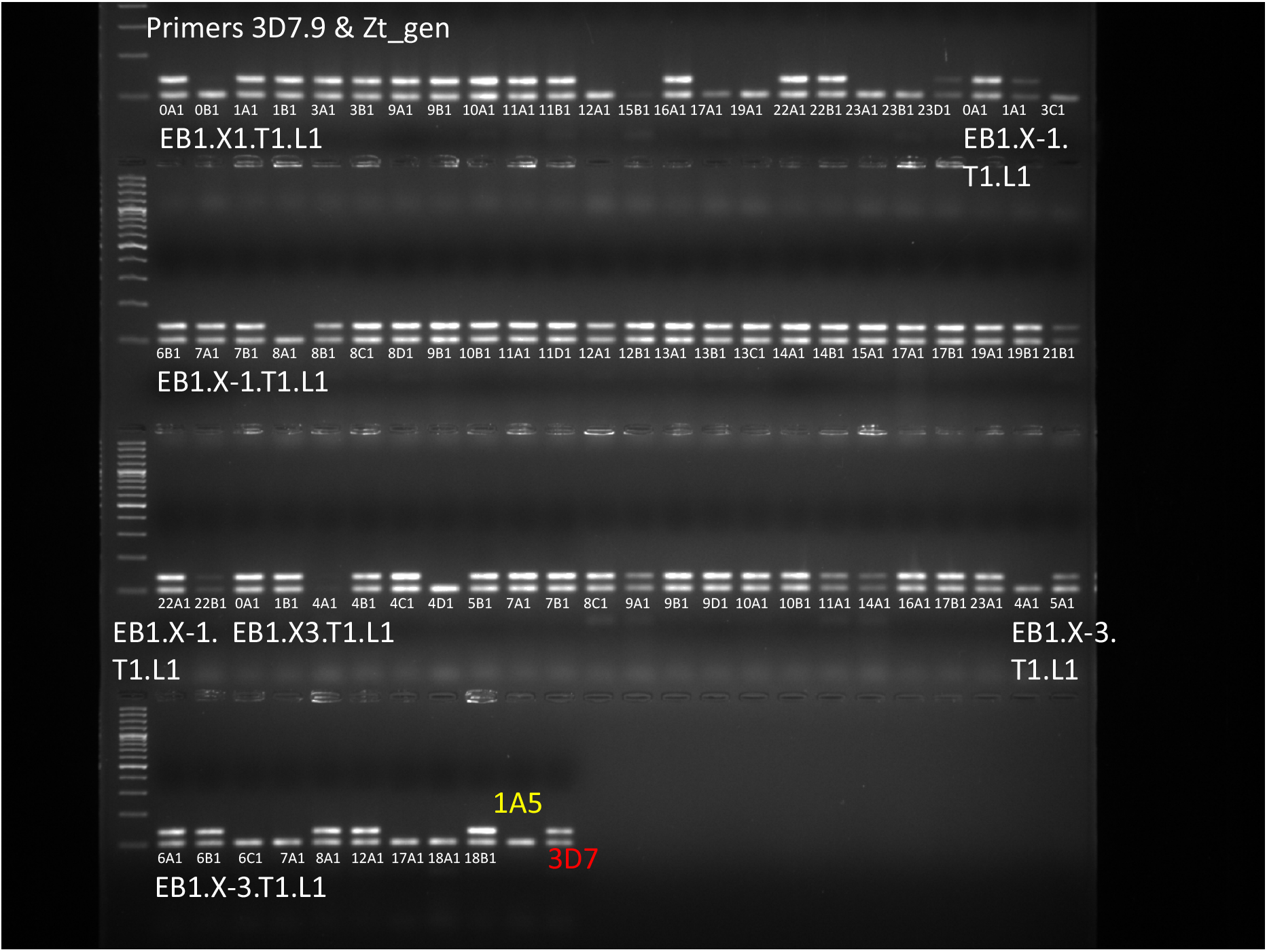
Amplicons from Zt_gen (shorter), 3D7.9 (longer). Isolate 3D7 as positive control and 1A5 as negative control. See Fig. A11 for label decoding.

**Figure A17:**
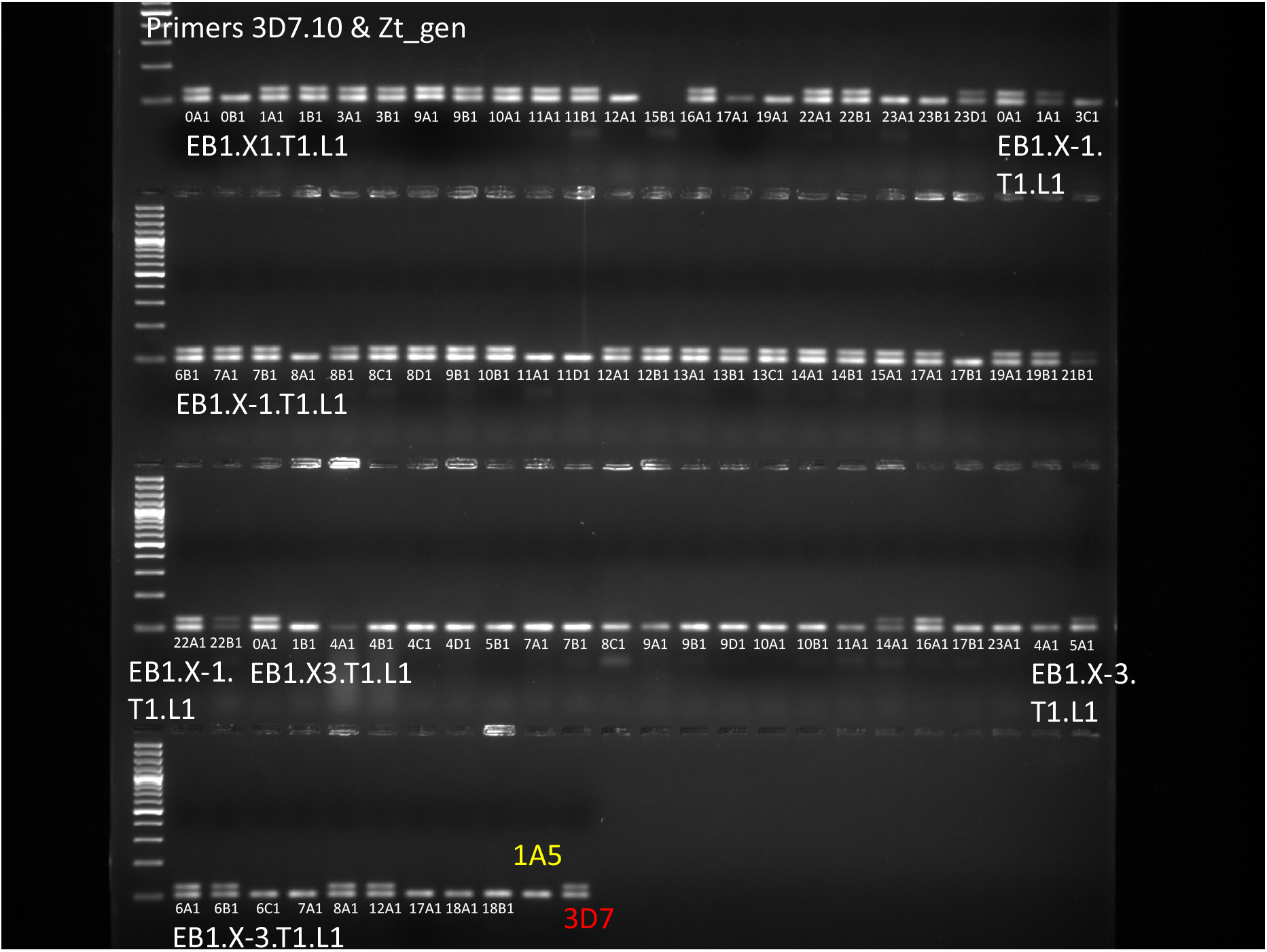
Amplicons from Zt_gen (shorter), 3D7.10 (longer). Isolate 3D7 as positive control and 1A5 as negative control. See Fig. A11 for label decoding.

## Appendix B: Pictures of inoculation

**Figure B1:**
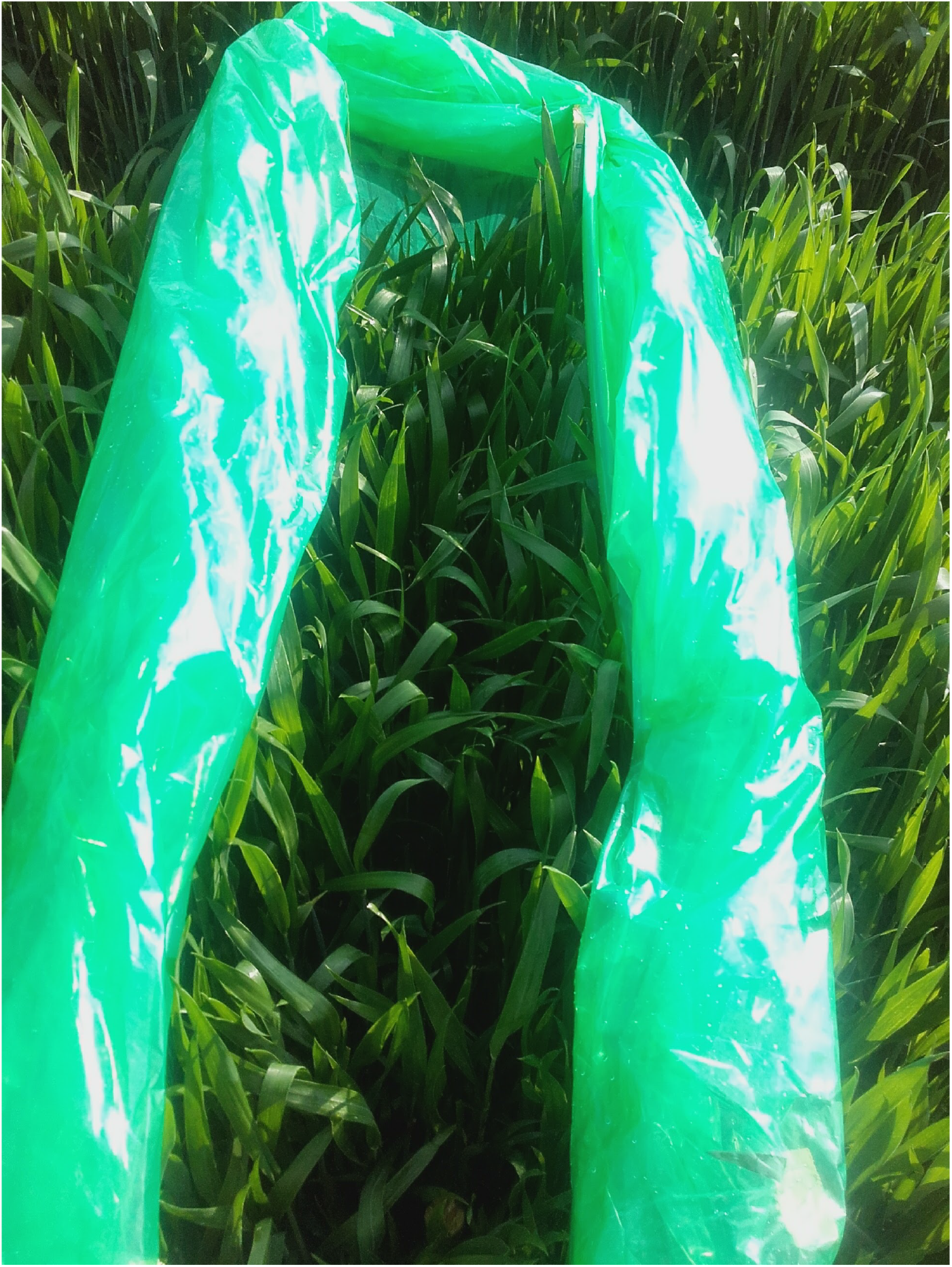
Inoculation tent prepared.

**Figure B2:**
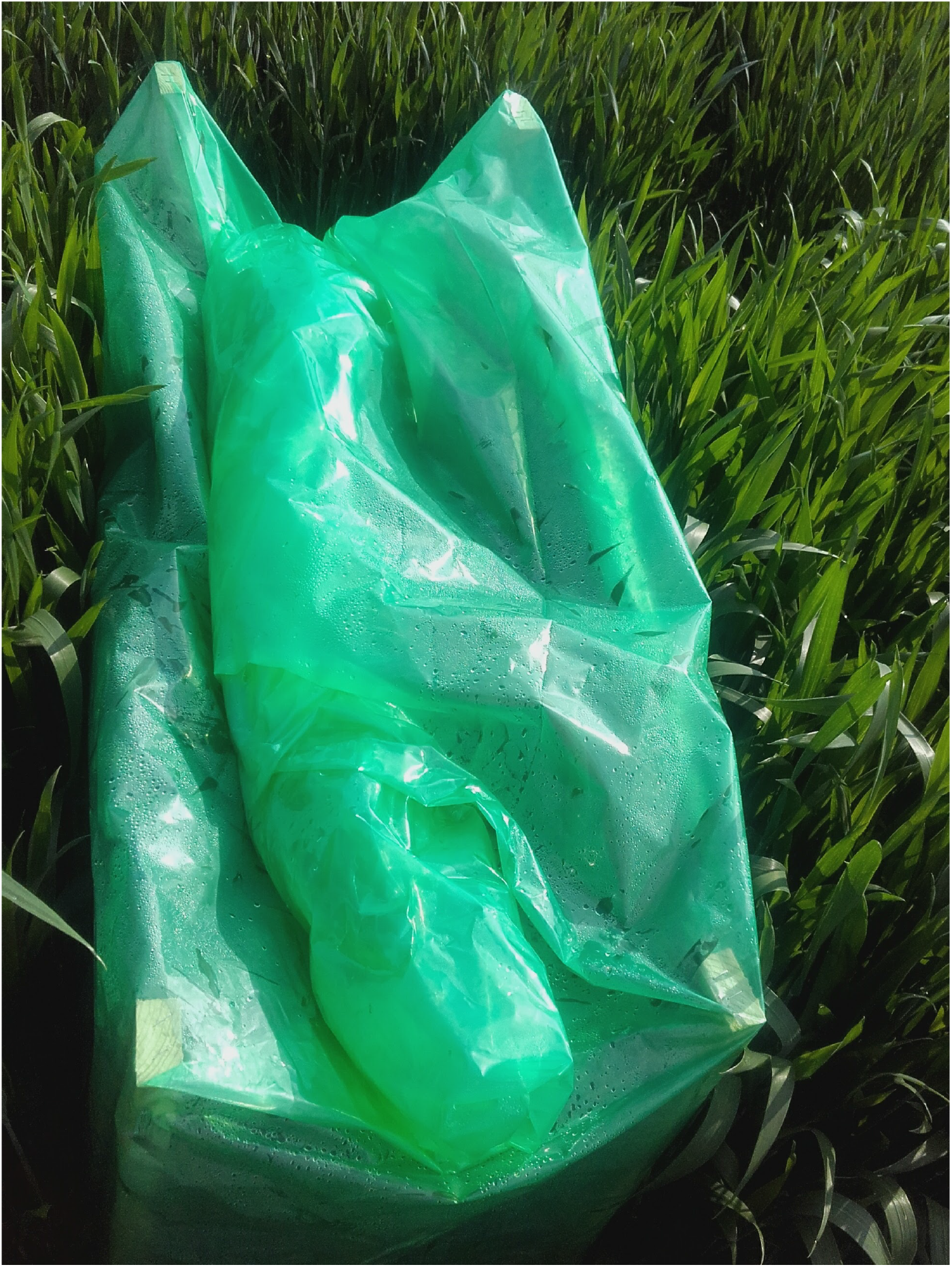
A tent closed after inoculation to maintain high humidity.

## Literature Cited

Anderegg, J., A. Hund, P. Karisto, and A. Mikaberidze. 2019. In-Field Detection and Quantification of Septoria Tritici Blotch in Diverse Wheat Germplasm Using Spectral-Temporal Features. Frontiers in Plant Science 10:1355.

Bannon, F., and B. Cooke. 1998. Studies on dispersal of Septoria tritici pycnidiospores in wheat-clover intercrops. Plant Pathology 47:49–56.

Ben M’Barek, S., P. Karisto, W. Abdedayem, M. Laribi, M. Fakhfakh, H. Kouki, A. Mikaberidze, and A. Yahyaoui. 2020. Improved control of septoria tritici blotch in durum wheat using cultivar mixtures. Plant Pathology 69:1655–1665.

Brennan, R., B. D. Fitt, G. Taylor, and J. Colhoun. 1985. Dispersal of Septoria nodorum pycnidiospores by simulated raindrops in still air. Journal of Phytopathology 112:281–290.

Brophy, L. S., and C. C. Mundt. 1991. Influence of plant spatial patterns on disease dynamics, plant competition and grain yield in genetically diverse wheat populations. Agriculture, ecosystems & environment 35:1–12.

Davison, A. C., and D. V. Hinkley. 1997. Bootstrap methods and their application. Cambridge university press.

Dean, R., J. A. van Kan, Z. A. Pretorius, K. E. Hammond-Kosack, A. Di Pietro, P. D. Spanu, J. J. Rudd, M. Dickman, R. Kahmann, J. Ellis, and G. D. Foster. 2012. The Top 10 fungal pathogens in molecular plant pathology. Molecular plant pathology 13:414–430.

Djidjou-Demasse, R., B. Moury, and F. Fabre. 2017. Mosaics often outperform pyramids: insights from a model comparing strategies for the deployment of plant resistance genes against viruses in agricultural landscapes. New Phytologist 216:239–253.

Duvivier, M., G. Dedeurwaerder, M. De Proft, J.-M. Moreau, and A. Legrève. 2013. Real-time PCR quantification and spatio-temporal distribution of airborne inoculum of Mycosphaerella graminicola in Belgium. European Journal of Plant Pathology 137:325–341.

Fitt, B. D., H. McCartney, and P. Walklate. 1989. The role of rain in dispersal of pathogen inoculum. Annual review of phytopathology 27:241–270.

Fitt, B. D. L., P. H. Gregory, A. D. Todd, H. A. McCartney, and O. C. Macdonald. 1987. Spore dispersal and plant disease gradients; a comparison between two empirical models. Journal of Phytopathology 118:227–242.

Gregory, P. 1968. Interpreting plant disease dispersal gradients. Annual Review of Phytopathology 6:189–212.

Hartmann, F. E., and D. Croll. 2017. Distinct trajectories of massive recent gene gains and losses in populations of a microbial eukaryotic pathogen. Molecular biology and evolution 34:2808–2822.

Hartmann, F. E., A. Sánchez-Vallet, B. A. McDonald, and D. Croll. 2017. A fungal wheat pathogen evolved host specialization by extensive chromosomal rearrangements. The ISME journal 11:1189–1204.

Heald, F. 1913. The Address of the President for 1912: The Dissemination of Fungi Causing Disease. Transactions of the American Microscopical Society 32:5–29.

Henze, M., M. Beyer, H. Klink, and J.-A. Verreet. 2007. Characterizing meteorological scenarios favorable for Septoria tritici infections in wheat and estimation of latent periods. Plant Disease 91:1445–1449.

Johansson, R., P. Strålfors, and G. Cedersund. 2014. Combining test statistics and models in bootstrapped model rejection: it is a balancing act. BMC systems biology 8:46.

Jørgensen, L. N., M. S. Hovmøller, J. G. Hansen, P. Lassen, B. Clark, R. Bayles, B. Rode-mann, K. Flath, M. Jahn, T. Goral, J. J. Czembor, P. Cheyron, C. Maumene, C. D. Pope, R. Ban, G. C. Nielsen, and G. Berg. 2014. IPM Strategies and Their Dilemmas Including an Introduction to www.eurowheat.org. Journal of Integrative Agriculture 13:265–281.

Karisto, P., S. Dora, and A. Mikaberidze. 2019a. Measurement of infection efficiency of a major wheat pathogen using time-resolved imaging of disease progress. Plant Pathology 68:163–172.

Karisto, P., A. Hund, K. Yu, J. Anderegg, A. Walter, F. Mascher, B. A. McDonald, and A. Mikaberidze. 2018. Ranking Quantitative Resistance to Septoria tritici Blotch in Elite Wheat Cultivars Using Automated Image Analysis. Phytopathology 108:568–581.

Karisto, P., F. Suffert, and A. Mikaberidze. 2019b. Spatially-explicit modeling improves empirical characterization of dispersal: theory and a case study. bioRxiv page 789156.

Kirchgessner, N., F. Liebisch, K. Yu, J. Pfeifer, M. Friedli, A. Hund, and A. Walter. 2017. The ETH field phenotyping platform FIP: a cable-suspended multi-sensor system. Functional Plant Biology 44:154–168.

Klein, E. K., A. Bontemps, and S. Oddou-Muratorio. 2013. Seed dispersal kernels estimated from genotypes of established seedlings: does density-dependent mortality matter? Methods in Ecology and Evolution 4:1059–1069.

Lovell, D., T. Hunter, S. Powers, S. Parker, and F. van den Bosch. 2004a. Effect of temperature on latent period of septoria leaf blotch on winter wheat under outdoor conditions. Plant Pathology 53:170–181.

Lovell, D., S. Parker, T. Hunter, D. Royle, and R. Coker. 1997. Influence of crop growth and structure on the risk of epidemics by Mycosphaerella graminicola (Septoria tritici) in winter wheat. Plant pathology 46:126–138.

Lovell, D., S. Parker, T. Hunter, S. Welham, and A. Nichols. 2004b. Position of inoculum in the canopy affects the risk of septoria tritici blotch epidemics in winter wheat. Plant Pathology 53:11–21.

Madden, L. V., G. Hughes, and F. van den Bosch. 2007. The study of plant disease epidemics. American Phytopathological Society (APS Press).

McCartney, H., B. D. Fitt, and J. S. West, 2006. Dispersal of foliar plant pathogens: mechanisms, gradients and spatial patterns. Pages 159–192 in B. M. Cooke, D. G. Jones, and B. Kaye, editors. The epidemiology of plant diseases. Springer.

Mikaberidze, A., B. A. McDonald, and S. Bonhoeffer. 2014. Can high-risk fungicides be used in mixtures without selecting for fungicide resistance? Phytopathology 104:324–331.

Mikaberidze, A., N. D. Paveley, S. Bonhoeffer, and F. van den Bosch. 2017. Emergence of fungicide resistance: the role of fungicide dose. Phytopathology 107:1–16.

Morais, D., V. Laval, I. Sache, and F. Suffert. 2015. Comparative pathogenicity of sexual and asexual spores of Zymoseptoria tritici (Septoria tritici blotch) on wheat leaves. Plant Pathology 64:1429–1439.

Morais, D., I. Sache, F. Suffert, and V. Laval. 2016. Is the onset of septoria tritici blotch epidemics related to the local pool of ascospores? Plant Pathology 65:250–260.

Mundt, C., and J. Browning. 1985. Development of crown rust epidemics in genetically diverse oat populations: effect of genotype unit area. Phytopathology 75:607–610.

Nathan, R., E. Klein, J. J. Robledo-Arnuncio, and E. Revilla, 2012. Dispersal kernels: review. Chapter 15, pages 187–210 in J. Clobert, M. Baguette, T. G. Benton, and J. M. Bullock, editors. Dispersal ecology and evolution. Oxford University Press Oxford.

Newton, A., G. Begg, and J. Swanston. 2009. Deployment of diversity for enhanced crop function. Annals of Applied Biology 154:309–322.

Newton, A., and D. Guy. 2011. Scale and spatial structure effects on the outcome of barley cultivar mixture trials for disease control. Field Crops Research 123:74–79.

Newville, M., T. Stensitzki, D. B. Allen, and A. Ingargiola. 2014. LMFIT: Non-Linear Least-Square Minimization and Curve-Fitting for Python. Zenodo.

Robert, C., G. Garin, M. Abichou, V. Houlès, C. Pradal, and C. Fournier. 2018. Plant architecture and foliar senescence impact the race between wheat growth and Zymoseptoria tritici epidemics. Annals of botany 121:975–989.

Saint-Jean, S., M. Chelle, and L. Huber. 2004. Modelling water transfer by rain-splash in a 3D canopy using Monte Carlo integration. Agricultural and Forest Meteorology 121:183–196.

Sapoukhina, N., Y. Tyutyunov, I. Sache, and R. Arditi. 2010. Spatially mixed crops to control the stratified dispersal of airborne fungal diseases. Ecological Modelling 221:2793–2800.

Shaw, M. 1987. Assessment of upward movement of rain splash using a fluorescent tracer method and its application to the epidemiology of cereal pathogens. Plant Pathology 36:201–213.

Shaw, M. 1990. Effects of temperature, leaf wetness and cultivar on the latent period of Mycosphaerella graminicola on winter wheat. Plant Pathology 39:255–268.

Shaw, M., and D. Royle. 1993. Factors determining the severity of epidemics of Mycosphaerella graminicola (Septoria tritici) on winter wheat in the UK. Plant Pathology 42:882–899.

Stewart, E. L., D. Croll, M. H. Lendenmann, A. Sanchez-Vallet, F. E. Hartmann, J. Palma-Guerrero, X. Ma, and B. A. Mcdonald. 2018. Quantitative trait locus mapping reveals complex genetic architecture of quantitative virulence in the wheat pathogen Zymoseptoria tritici. Molecular plant pathology 19:201–216.

Suffert, F., G. Delestre, and S. Gélisse. 2018. Sexual Reproduction in the Fungal Foliar Pathogen Zymoseptoria tritici Is Driven by Antagonistic Density Dependence Mechanisms. Microbial ecology 77:1–14.

Suffert, F., and R. N. Thompson. 2018. Some reasons why the latent period should not always be considered constant over the course of a plant disease epidemic. Plant Pathology 67:1831–1840.

Terpilowski, M., 2018. Scikit-posthocs. URL https://scikit-posthocs.readthedocs.io.

Vidal, T., C. Gigot, C. de Vallavieille-Pope, L. Huber, and S. Saint-Jean. 2018. Contrasting plant height can improve the control of rain-borne diseases in wheat cultivar mixture: modelling splash dispersal in 3-D canopies. Annals of botany 121:1299–1308.

Vidal, T., P. Lusley, M. Leconte, C. De Vallavieille-Pope, L. Huber, and S. Saint-Jean. 2017. Cultivar architecture modulates spore dispersal by rain splash: A new perspective to reduce disease progression in cultivar mixtures. PloS one 12:e0187788.

Virtanen, P., R. Gommers, T. E. Oliphant, M. Haberland, T. Reddy, D. Cournapeau, E. Burovski, P. Peterson, W. Weckesser, J. Bright, S. J. van der Walt, M. Brett, J. Wilson, K. Jarrod Millman, N. Mayorov, A. R. J. Nelson, E. Jones, R. Kern, E. Larson, C. Carey, İ. Polat, Y. Feng, E. W. Moore, J. Vand erPlas, D. Laxalde, J. Perktold, R. Cimrman, I. Henriksen, E. A. Quintero, C. R. Harris, A. M. Archibald, A. H. Ribeiro, F. Pedregosa, P. van Mulbregt, and SciPy 1. 0 Contributors. 2020. SciPy 1.0: Fundamental Algorithms for Scientific Computing in Python. Nature Methods 17:261–272.

Willocquet, L., S. Savary, B. A. McDonald, and A. Mikaberidze. 2020. A polyetic modelling framework for plant disease emergence. Plant Pathology 69:1630–1643.

Yu, K., J. Anderegg, A. Mikaberidze, P. Karisto, F. Mascher, B. A. McDonald, A. Walter, and A. Hund. 2018. Hyperspectral canopy sensing of wheat septoria tritici blotch disease. Frontiers in plant science 9:1195.

Zadoks, J., and F. van den Bosch. 1994. On the spread of plant disease: a theory on foci. Annual review of phytopathology 32:503–521.

Zadoks, J. C., T. T. Chang, and C. F. Konzak. 1974. A decimal code for the growth stages of cereals. Weed research 14:415–421.

Zhan, J., G. H. Kema, C. Waalwijk, and B. A. McDonald. 2002. Distribution of mating type alleles in the wheat pathogen Mycosphaerella graminicola over spatial scales from lesions to continents. Fungal Genetics and Biology 36:128–136.

Zhan, J., C. Mundt, and B. McDonald. 1998. Measuring immigration and sexual reproduction in field populations of Mycosphaerella graminicola. Phytopathology 88:1330–1337.

Zhan, J., C. Mundt, and B. McDonald. 2000. Estimation of rates of recombination and migration in populations of plant pathogens—a reply. Phytopathology 90:324–326.

